# An unconventional GABAergic circuit differently controls pyramidal neuron activity in two visual cortical areas via endocannabinoids

**DOI:** 10.1101/2021.09.06.459113

**Authors:** Martin Montmerle, Fani Koukouli, Andrea Aguirre, Jérémy Peixoto, Vikash Choudhary, Marcel De Brito Van Velze, Marjorie Varilh, Francisca Julio-Kalajzic, Camille Allene, Pablo Mendéz, Giovanni Marsicano, Oliver M. Schlüter, Nelson Rebola, Alberto Bacci, Joana Lourenço

## Abstract

Perisomatic inhibition of neocortical pyramidal neurons (PNs) coordinates cortical network activity during sensory processing, and it has been mainly attributed to parvalbumin-expressing basket cells (BCs). However, cannabinoid receptor type 1 (CB1)-expressing interneurons also inhibit the perisomatic region of PNs but the connectivity and function of these elusive – yet prominent – neocortical GABAergic cells is unknown. We found that the connectivity pattern of CB1-positive BCs strongly differs between primary and high-order cortical visual areas. Moreover, persistently active CB1 signaling suppresses GABA release from CB1 BCs in the medial secondary visual cortex (V2M), but not in the primary (V1) visual area. Accordingly, *in vivo*, tonic CB1 signaling is responsible for higher but less coordinated PN activity in V2M than in V1. Our results indicate a differential CB1-mediated mechanism controlling PN activity, and suggest an alternative connectivity schemes of a specific GABAergic circuit in different cortical areas

## Introduction

The integration of sensory information into a coherent representation of the external world is accomplished by cortical circuits formed by a multitude of cellular subtypes that connect with each other following a detailed blueprint (Pfeffer et al., 2013;Kepecs and Fishell, 2014;Allene et al., 2015;Tremblay et al., 2016). Importantly, fast synaptic GABAergic inhibition is crucial in shaping both spontaneous and sensory-evoked cortical activity (Isaacson and Scanziani, 2011). Inhibitory GABAergic neurons (interneurons) are highly heterogeneous, and this remarkably rich diversity of cell types is epitomized by the specific pattern of synaptic connections that each interneuron subtype forms with excitatory principal neurons (PNs) (Ascoli et al., 2008;Tremblay et al., 2016;Lourenco et al., 2020b). The result is an efficient orchestration of cortical activity (Isaacson and Scanziani, 2011) *via* a highly specialized division of labor of different interneuron types (Freund and Katona, 2007). In particular, perisomatic-targeting basket cells (BCs) form inhibitory synapses mainly near the cell soma of PNs, and thus control spike generation and timing (Buzsaki and Wang, 2012). A prominent perisomatic-targeting interneuron type, the parvalbumin (PV)-expressing BC, is important during sensory information processing (Atallah et al., 2012;Lee et al., 2012;Wilson et al., 2012). Moreover, due to their fast biophysical kinetics and reliability, PV BCs drive cognitive-relevant network oscillations in the γ-frequency range (30-100 Hz) (Freund and Katona, 2007;Buzsaki and Wang, 2012;Deleuze et al., 2019). PV BCs have been extensively studied because they are abundant, easily identifiable by their typical fast-spiking pattern, and they can be genetically tagged and manipulated in different brain regions (Taniguchi et al., 2011). Yet, PV BCs are not the only inhibitory cell type controlling the perisomatic region of neocortical PNs (Freund and Katona, 2007). Most notably, interneurons expressing high levels of cannabinoid receptor type 1 (CB1) also form inhibitory synapses with PN cell bodies (Bodor et al., 2005). These cells have been traditionally identified as expressing cholecystokinin (CCK), especially in the hippocampus (Marsicano and Lutz, 1999;Katona et al., 1999;Hefft and Jonas, 2005;Daw et al., 2009;Dudok et al., 2021), and they form GABAergic synapses that are much less reliable (Wilson et al., 2001;Hefft and Jonas, 2005;Daw et al., 2009). However, the properties and roles of CB1 BCs within neocortical circuits remain elusive. In particular, whether CB1 BCs operate an efficient control of neocortical PN firing is unknown.

Endogenous cannabinoids (endocannabinoids, eCBs) acting on CB1 are known to modulate strongly synaptic transmission (Hajos et al., 2000;Maejima et al., 2001;Wilson and Nicoll, 2001;Kano et al., 2009;Castillo et al., 2012). In particular, CB1 receptors potently inhibit release of GABA, resulting in several forms of inhibitory plasticity, such as depolarization-induced suppression of inhibition (DSI) and long-term depression of inhibitory synapses (LTDi) (Marsicano et al., 2002;Younts and Castillo, 2014). Both plastic phenomena rely on retrograde signaling of eCBs, which are synthesized on demand in the postsynaptic pyramidal neurons (PNs) by intracellular Ca^2+^ increase or mGluR activation, and delivered to presynaptic terminals of CB1-expressing GABAergic interneurons (Kano et al., 2009;Castillo et al., 2012). In addition to this on-demand eCB modulation of neurotransmitter release, in the hippocampus, CB1 receptors were reported to be persistently active thus leading to constant signaling and tonic inhibition of GABA release from CB1 interneurons (Losonczy et al., 2004;Neu et al., 2007;Lee et al., 2010).

Sensory systems are highly organized and hierarchal. Sensory information is relayed (via the thalamus) to the primary sensory neocortical areas, mainly in layer (L) 4, and it is then passed along to other cortical layers in a stereotyped sequence before being sent to higher-order associative cortical areas. In parallel, higher-order cortical areas send information to primary cortices, modulating their activity (Larkum, 2013). These loops allow information to travel across different sensory areas via both direct connections and cortico-thalamic pathways (Larkum, 2013;Harris and Shepherd, 2015;Glickfeld and Olsen, 2017). Higher-order sensory cortices are integrative areas receiving both bottom-up (or sensory) as well as top-down (or contextual) information, thus playing a major role in decoding specific sensory features and in predictive processing and behavior (Glickfeld and Olsen, 2017;Clancy et al., 2019;Keller et al., 2020;Murgas et al., 2020;Jin and Glickfeld, 2020). Cognitive activity is strongly influenced by the functional state of cortical circuits, which have been hypothesized to be uniform across different cortical areas and characterized by specific functions (Kepecs and Fishell, 2014). Nevertheless, recent evidence indicates that the canonical cortical circuit for perisomatic inhibition exhibits peculiarities in different cortical areas (Whissell et al., 2015;Pluta et al., 2019;Naka et al., 2019;Lourenco et al., 2020b).

Using a mouse line specifically tagging CB1-expressing neurons (CB1-tdTomato mice) (Winters et al., 2012), here we set out to study the morpho-functional features of neocortical CB1 BCs and test whether these cells operate an efficient control of PN activity in the visual cortex.

In contrast to generally thought homogenous interneuron motifs, we found that CB1 expression is generally stronger in associative than in primary sensory cortical areas. We then focused on primary visual cortex (V1) and its associative, higher-order medial secondary area (V2M), which exhibits strong CB1 expression. We describe a differential morphological and functional connectivity scheme of CB1 interneurons in V1 and V2M. Tonic CB1 modulation conferred specific weak presynaptic properties at inhibitory synapses from CB1 BCs only in L2/3 of V2M. This area-specific connectivity and eCB modulation of GABA release from CB1 BCs was responsible for lower PN activity in V1 compared to V2M *in vivo*. Moreover, visual area-specific tonic CB1 signaling differently set the amount of correlated activity of PNs in the two visual areas. Therefore, tonic CB1 signaling is responsible for a dynamic modulation of GABA release, which is specific for higher order visual areas.

## Results

### Differential CB1 expression and morphological properties of CB1-postitive interneurons in primary and higher-order visual cortex

CB1 is known to modulate neurotransmitter release in virtually all brain areas (Castillo et al., 2012). In particular, CB1-dependent modulation of GABA release has been well characterized in hippocampal interneurons, but whether neocortical CB1-expressing interneurons possess similar properties is unclear. Moreover, differences in CB1 expression between primary and higher-order areas have been reported (Yoneda et al., 2013) and could account for functional diversity of GABAergic circuits across different cortical areas. However, the exact CB1 expression pattern across layers in V1 and V2M has not yet been examined. We therefore sought to describe this pattern in more detail using CB1 immunofluorescence in wild-type mice (Figure 1A). CB1 immunostaining revealed a classical pattern of fibrous processes, which is consistent with its main axonal expression. Overall, CB1 expression was highest in L1 of V1 and in L2/3 of V2M (Figure 1A and 1B). To compare the laminar expression between the two areas, we binned this data into distances corresponding to cortical layers (Figure 1C). In V1, the average intensity of peak-normalized fluorescence gradually decreased from L1 to L4 (respectively 70.5±2.2 % and 38.7±2.3 %, n=11 animals, 3 slices per animal), to then increase and reach another peak in L6 (66.3±3.2 %; Figure 1A, black and 1B, black bars). In contrast, in V2M the peak-normalized fluorescence from pia to white matter was always comprised between a maximum in L2/3 and a minimum in L5 (respectively 70.7±3.3 % and 57.3±3.3 %; Figure 1A, red line and 1B, red bars). We found that, except L6 and L1, the level of expression of CB1 was higher in all other cortical layers of V2M than V1 (Friedman, repeated-measures, post hoc analysis with correction for multiple comparisons, Sidak’s multiple comparisons test; Figure 1B,C). In particular, L2/3, 4 and 5 of V1 CB1 expression was weaker than in V2M (52.6 ± 2.2, 38.77 ± 2.3, 41 ± 2.2%, respectively; p=0.0001, p<0.0001, p= 0.0006, respectively).

**Figure 1.**
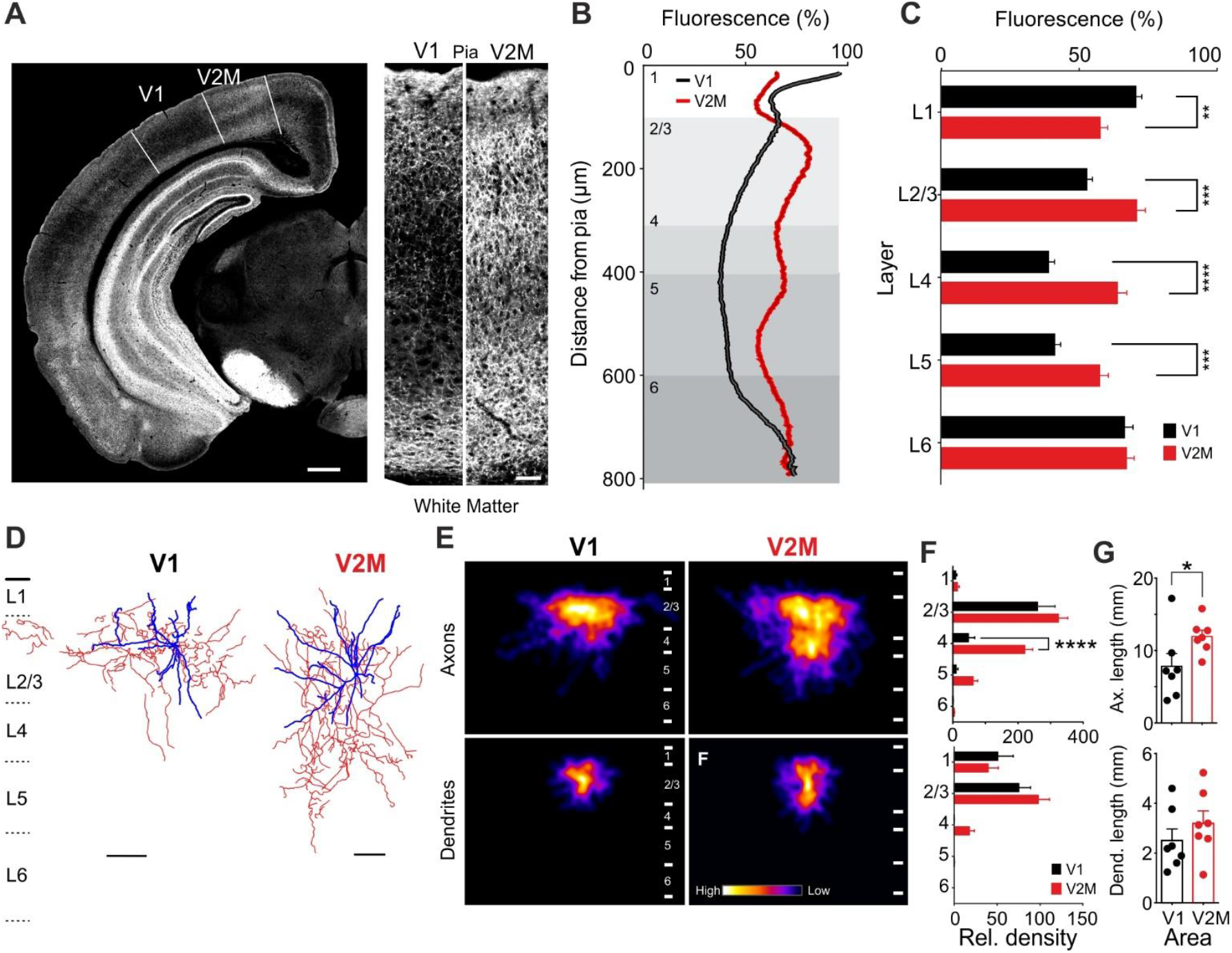
Differential CB1 expression and morphological properties of CB1 interneurons in primary and secondary visual cortex. **A:** Left: Micrograph illustrating CB1 immunofluorescence in a coronal slice of a mouse brain including V1 and V2M (delimited by dashed lines). Scale: 500 µm. Right: Blowout and side-by-side comparison of CB1 immunofluorescence between V1 and V2M of the same slice, using the identical acquisition settings. Scale: 50 µm. **B:** CB1 fluorescence intensity measured from pia to white matter in V1 and V2M, normalized to the peak of fluorescence in both region. Layers are delimited by white/grey boxes, with numbers in top left corners. **C:** Data from B binned into cortical layer. Error bars are SEM. **p<0.01 and****p<0.0001; n=3 slices per animal, N = 11 animals. **D:** Representative reconstructed CB1 BCs in V1 (left) and V2M (right). Red and blue tracing refer to axons and dendrites, respectively. Scale bar: 100 µm. **E:** Heat map of CB1 BC axonal density in V1 (left) and V2M (right). Numbers designate cortical layers. All filled neurons (n = 7, both areas) were overlapped to generate this figure. **F:** Relative axonal (top) and dendritic (bottom) density, normalized by layer thickness (see methods). *p<0.05 and **p<0.01. **G**: Total axonal (top) and dendritic (bottom) length in the two areas.

CB1 is most prominently expressed on inhibitory axonal fibers (Marsicano and Lutz, 1999;Katona et al., 1999). Hence, the difference in CB1 expression pattern between V1 and V2M shown in Figure 1 could be explained by: *i)* different numbers of CB1 BCs between the two areas, *ii)* distinct fiber density originating from the same interneurons, *iii)* different level of expression of CB1 in each given axon. CB1 is not expressed exclusively by GABAergic interneurons (Marsicano and Lutz, 1999;Katona et al., 2006;Marinelli et al., 2009). Therefore, we crossed CB1-tdTomato with GAD67-GFP mice, to quantify CB1-expressing GABAergic cells in the two visual areas. We thus counted CB1-positive GABAergic interneurons in each layer of the two areas (Figure S1). CB1-expressing interneurons were prominently expressed in L2/3, comprising 31% and 35% of the total CB1-IN population in V1 and V2M, respectively (Figure S1A). In particular, we found that the large majority of CB1 cells co-expressed GAD67 (Figure S1A; orange arrows; 85% and 93% co-localization in L2/3 V1 and V2M, respectively). We found that in L2/3 of V2M, CB1 interneurons were significantly more numerous than in V1 (Friedman, repeated-measures, post hoc analysis with correction for multiple comparisons, Sidak’s multiple comparisons test p<0.05; Figure S1C). Importantly, CB1 cells did not co-localize with parvalbumin (PV)- nor somatostatin (SST)-positive cells (Figure S2A,B), indicating that CB1 BCs do not belong to these two prominent cortical interneuron subclasses originating from the medial ganglionic eminence (MGE). However, the majority (∼60%) of CB1 cells co-localized with 5HT_3A_R (Figure S2 C,D). In situ hybridization using *CB_1_* (*Cnr1*) and *CCK (Cck)* mRNA probes revealed co-localization of td-Tomato expressing neurons with *CB1* (>75%) and *CCK* (>40%; Figure S2 G). Overall, these results indicate that the CB1-tdTomato accurately labels CB1 neurons, which are part of interneurons originating from the caudal ganglionic eminence (CGE) (Tremblay et al., 2016;Paul et al., 2017).

To test whether increased CB1 staining in deeper cortical layers in V2M was also due to differential axonal projections, we performed whole-cell, patch-clamp recordings from L2/3 multipolar CB1 BCs, visually identified as expressing bright fluorescence in CB1-tdTomato mice. We included high concentrations of biocytin (5-8 mg/mL) in whole-cell pipettes to reveal axonal arborizations (Jiang et al., 2015). Untypical for local soma innervating interneurons, anatomically reconstructed CB1 BCs revealed a strong axonal innervation in deep cortical L4 selectively in V2M, whereas the same type of neurons in V1 more typically projected mainly within L2/3 (Figure 1 D-F and Figure S3; relative axonal density: 220 ± 25 and 62 ± 15 vs. 47 ± 20 and 10 ± 6, V2M vs. V1, respectively; p < 0.001; n = 7 for both V1 and V2M). Accordingly, CB1 BCs in V2M exhibited a larger total axonal length than in V1 (p= 0.0379; Mann–Whitney U test), with no differences in dendritic length in both visual areas (p = 0.32; Mann– Whitney U test; Figure 1G).

Differences in axonal projection patterns between CB1 BCs in V1 and V2M could imply that they belong to different functional interneuron subtypes. We therefore tested a battery of electrophysiological parameters, and found no difference in single action potential waveform and firing dynamics (Figure S4).

Altogether, these results indicate a contrasting pattern of CB1 expression in V1 and V2M. This is due to a more prominent descending axonal innervation selectively in V2M, suggesting a differential connectivity scheme from this IN subtype in the two visual areas.

### Functional differences of intra- and infra-layer connectivity of CB1 BCs in V1 and V2M

The morphological differences between L2/3 CB1 BCs in V1 and V2M described above prompted the question whether the probability of CB1 cells to connect to postsynaptic targets within and across layers is different in V1 and V2M.

We performed dual simultaneous whole-cell paired recordings between presynaptic CB1 BCs in L2/3 and postsynaptic PNs in either L2/3 or L4, in V1 and V2M (Figure 2A,B). In both areas, we reliably obtained connected pairs when the post-synaptic PN was also located in L2/3 (Figure 2A; connected pairs: 56 out of 218 vs. 53 out of 169 in V1 vs. V2M). However, when the postsynaptic cell was located in L4, the likelihood of obtaining connected pairs was very low in V1 while remaining high in V2M (5 out of 75 and 46 out of 187, respectively; Figure 2A). These connectivity rates are largely consistent with the axonal morphologies of these cells (Figure 1D-G). We found that the amplitudes of unitary IPSCs (uIPSCs) were ∼5.5 fold larger onto PNs in L2/3 of V1 than V2M (uIPSC amplitude : 183.4 ± 39.93 pA vs. 33.38± 5.539 pA, L2/3 V1 vs. L2/3 V2M, n = 44 and 40, respectively, p<0.0001; Kurskal-Wallis

**Figure 2.**
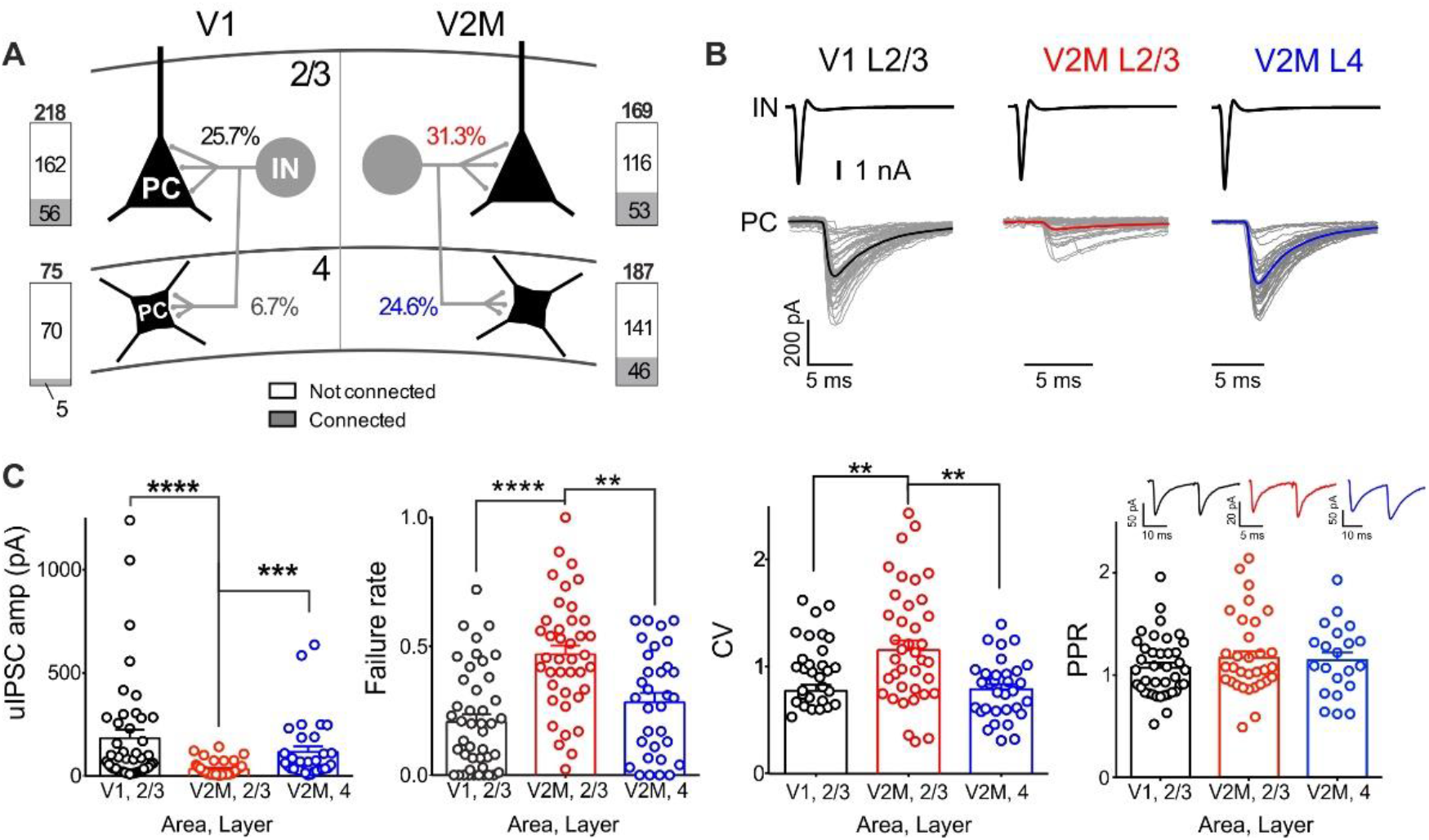
Functional differences of intra- and infra-layer connectivity of CB1 BCs in V1 and V2M. **A**: Connectivity rates of CB1 BC → PN connectivity within L2/3 and to L4 in both V1 and V2M. **B:** Representative voltage-clamp traces of evoked uIPSCs in the postsynaptic PN (presynaptic action current above, in black) in monosynaptically connected pairs in V1 L2/3 (avg. in black), V2M L2/3 (avg. in red) and V2M L4 (avg. in blue) principal neurons. Individual (30) sweeps in light grey. **C:** Population data of uIPSC amplitude, failure rate, coefficient of variation (CV) and paired-pulse ratio (PPR). Black circles: V1 L2/3 connections; red circles: V2 L2/3 connections; blue circles: V2 L4 connections. All bar graphs represent mean ± SEM; *p<0.05, ****p<0.0001.

ANOVA followed by Dunn’s multiple comparisons test; Figure 2B and C). Accordingly, failure rate and variability (measured as coefficient of variation, or CV) of uIPSCs was much lower in CB1 IN-PN connections in L2/3 of V1 than V2M (p<0.0001 and p = 0.0028, respectively; Kurskal-Wallis ANOVA followed by Dunn’s multiple comparisons test; Figure 2C). When limiting the analysis to successes only, we did not observe any differences in the paired pulse ratio (PPR; p = 0.5, Kurskal-Wallis ANOVA; Figure 2C). Connected pairs between CB1 BCs and postsynaptic PNs located in L4 of V2M were stronger and more reliable than connections formed in L2/3 by the same interneurons (L4 uIPSCs amplitude : 116.4 ± 26.55pA, n = 32, p=0.0013, Kurskal-Wallis ANOVA followed by Dunn’s multiple comparisons test, Figure 2C). Indeed these infra-laminar connections (from L2/3 to L4 in V2M) had similar amplitudes, failure rate and CV as of CB1 IN-PN pairs in L2/3 of V1 (p>0.05, Kurskal-Wallis ANOVA for amplitude, failure rate and CV, Figure 2C). In sum, connections in L2/3 of V2M were the weakest and most unreliable, when compared to the neighboring area or layer. When we examined uIPSC rise times, we found that they were fast (<1ms) and similar for all three synaptic connections (Figure S5A,B). This is consistent with a perisomatic pattern of innervation, typical of basket cells (Figure S5C).

Overall, these data indicate remarkably different functional projection patterns in the two visual cortical areas. Moreover, inputs from CB1 BCs in the visual cortex have area- and layer-specific properties. Given the marked difference in CV and failure rate, it is likely that inhibitory synapses from these interneurons possess visual-area, layer- and target-specific differences of presynaptic release of GABA.

### Activity-dependent modulation of synaptic efficacy from CB1 BCs is visual area- and layer-specific

Activation of presynaptic CB1 is often followed by strong short- and long-term forms of plasticity (Castillo et al., 2012). One of the most prominently studied form of short-term (seconds) of GABAergic plasticity is depolarization-induced suppression of inhibition (DSI). We therefore tested whether CB1- mediated DSI was different depending on the visual cortical areas where uIPSCs were recorded. DSI was evoked by 5-s long postsynaptic depolarizations at 0 mV. We found that CB1-dependent DSI was robustly present in all connected pairs, and it similarly affected uIPSCs in both V1 and V2M and in both cortical layers (Figure S6). CB1 signaling can be modulated by presynaptic activity (Lourenco et al., 2014). Importantly, presynaptic GABA release can be persistently modulated by tonic activation of CB1 (Losonczy et al., 2004; Foldy et al., 2006; Neu et al., 2007), inducing chronic suppression of inhibitory responses, which are unreliable and weak, similarly to those exhibited by CB1 BCs targeting PNs in L2/3 of V2M. If this is the case, high frequency AP train invading the presynaptic terminal should be able to override this CB1-mediated tonic modulation of inhibitory transmission (Losonczy et al., 2004; Foldy et al., 2006; Neu et al., 2007). We therefore delivered four 50 Hz-trains to presynaptic CB1 BCs and found an up to two fold increase of synaptic efficacy (measured as total unitary postsynaptic charge) during train applications only at connections in V2M between CB1 interneurons and L2/3 PNs (mean synaptic charge: 2.5 ± 0.6 nC vs. 5.1 ± 0.97 nC train 1 vs. train 4; n = 16; p < 0.001, Wilcoxon matched-pairs signed rank test; Figure 3A-C). In contrast, in pairs between CB1 BCs and PNs in V1, L2/3 and V2M, L4 synaptic efficacy was similar across different presynaptic spike trains (mean synaptic charge: 6.6 ± 2.3 nC vs. 6.7 ± 2.3 nC train 1 vs. train 4 for V1, L2/3 and 6.6 ± 2.4 nC vs. 5.9 ± 1.5 nC train 1 vs. train 4 for V2M, L4; n =13 and n = 15, respectively; p>0.05, Wilcoxon matched-pairs signed rank test; Figure 3A-C).

**Figure 3.**
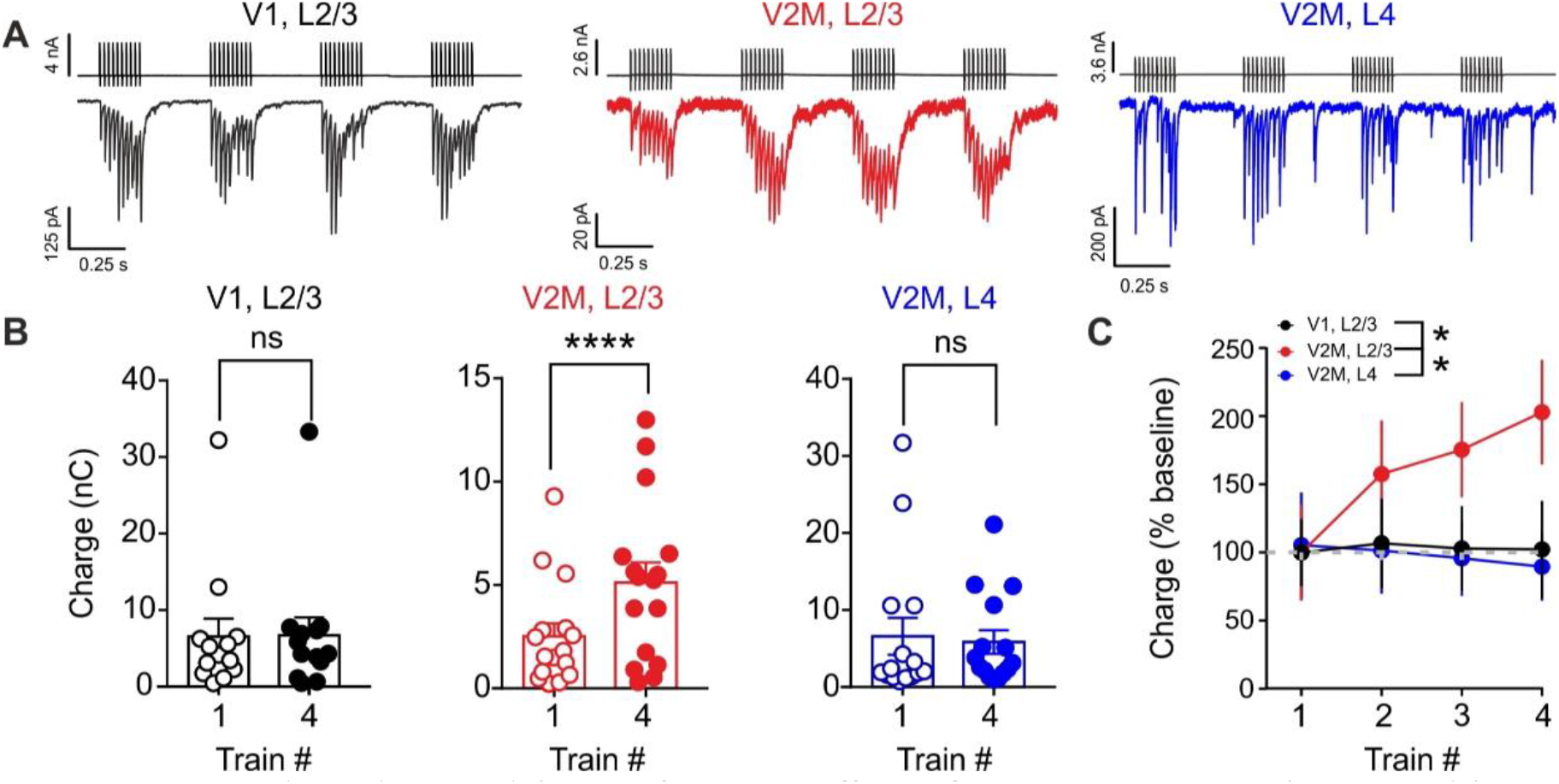
Activity-dependent modulation of synaptic efficacy from CB1 BCs is visual area- and layer- specific. **A:** Representative traces illustrating uIPSCs recorded in PNs, in response to four trains of presynaptic APs (10 spikes, 50 Hz) from CB1 BCs in V1, L2/3 (black), V2M, L2/3 (red) and V2M, L4 (blue). Shown are averaged traces of at least 10 individual trials. **B:** Population data of total postsynaptic charge calculated in the first and fourth train. **C:** Population data of percentage change of total postsynaptic charge at the three synapses in the two visual areas. Color code of B-C as in A.

These results indicate that GABAergic synapses contacting PNs in L2/3 of V2M were muffled selectively and persistently, but could be temporarily awakened by increased presynaptic activity to a level similar of the other studied connections.

### Visual area- and layer-specific GABAergic synaptic strength from CB1 BCs is due to selective tonic endocannabinoid signaling

Results of Figure 3 indicate that the low synaptic efficacy is specific for L2/3 connections between CB1 BCs and PNs in V2M. This suggest that tonic CB1 signaling is specific for this layer and visual area. We therefore, tested whether tonic signaling could play a role in explaining why the inhibitory synapses in L2/3 exhibit high failure rates and low amplitudes. We performed paired recordings in slices pre-incubated with the CB1 antagonist AM-251 (3 µM; Figure 4). In support of tonic CB1 signaling, the uIPSC amplitude between connected pairs in V2M – L2/3 increased, when AM-251 was present in the superfusate (uIPSC amplitude: 33 ± 6.3pA vs. 148 ± 51pA, n = 31 vs. 15 control vs. AM-251; p = 0.0254, Kurskal-Wallis ANOVA followed by Dunn’s multiple comparisons test, see below for comparisons with other synapses, Figure 4B), and the failure rate decreased (0.46 ± 0.04 vs. 0.25 ± 0.06, n = 31 vs. 15 control vs. AM-251; p = 0.0059, Mann Whitney test). Conversely, inter-stimulus variability (1.2 ± 0.1 vs. 0.87 ± 0.1, n = 31 vs. 15 control vs. AM-251; p = 0.0737, Mann Whitney test) was not significantly affected by AM-251 treatment. Overall, our data revealed that antagonizing CB1 converted weak and unreliable connections between CB1 BCs and PNs in L2/3 of V2M into strong and reliable synapses. Indeed, in the presence of AM-251, responses elicited in L2/3 of V2M were similar to those evoked in L2/3 of V1 and L4 of V2M (for both synapses p > 0.99 test, Kurskal-Wallis ANOVA followed by Dunn’s multiple comparisons test). In support of a selective effect in V2M L2/3, the drug had no effect on uIPSCs in V1, L2/3 and V2M, L4 (V1, L2/3 uIPSC amplitude: 208 ± 46 pA vs. 169 ± 49pA, n = 37 vs. 17 control vs. AM-251; V2M, L4 uIPSC amplitude: 133 ± 37 pA vs. 67 ± 13pA, n = 22 vs. 12 control vs. AM- 251; p>0.099 Kurskal-Wallis ANOVA followed by Dunn’s multiple comparisons test)

**Figure 4.**
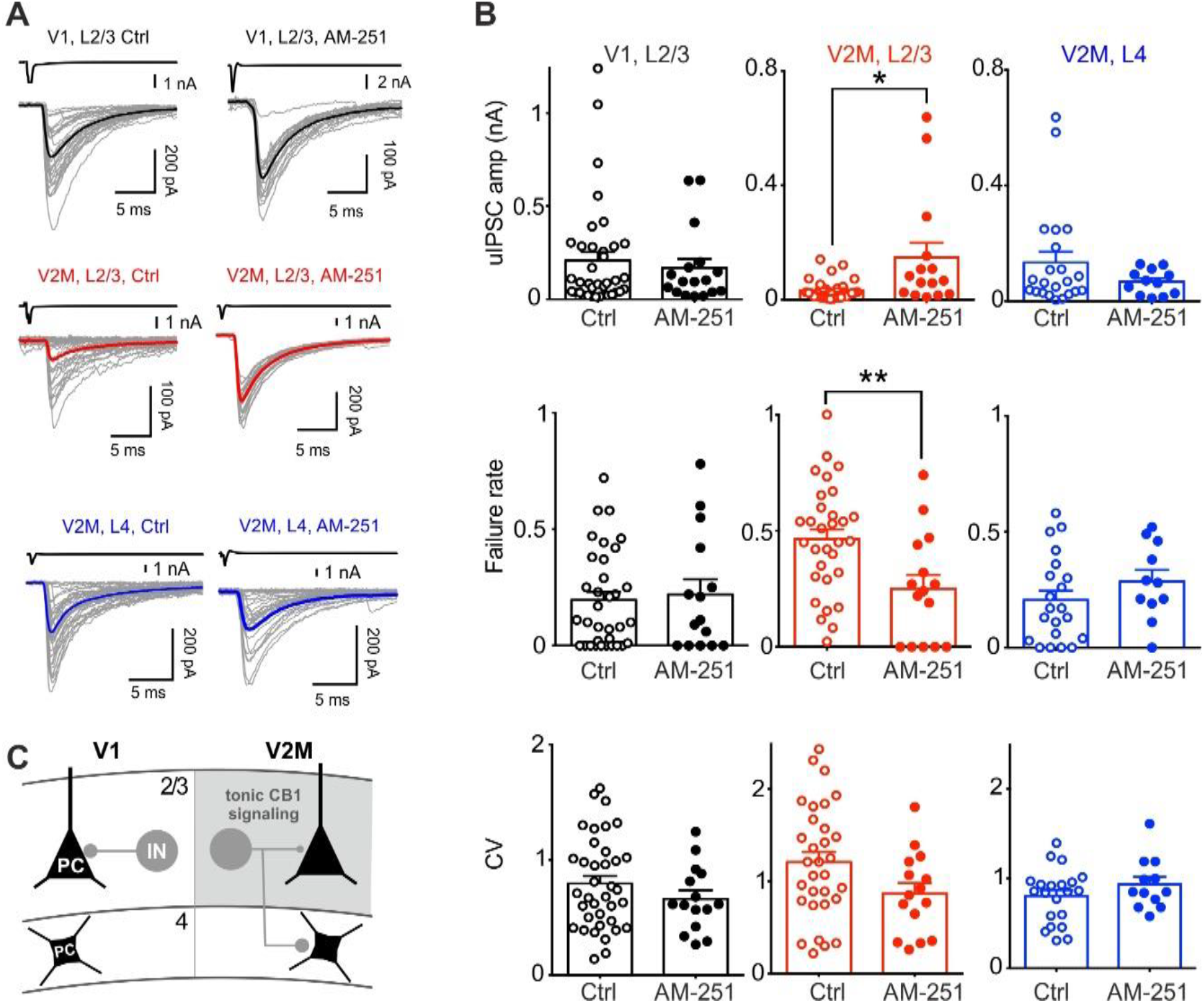
Visual area- and layer-specific GABAergic synaptic strength from CB1 BCs is due to selective tonic endocannabinoid signaling. **A:** Representative uIPSC traces in the absence (Ctrl, left) and presence of 3 µM AM-251 (right) in L2/3 of V1 (top, black), L2/3 of V2M (middle, red) and L4 of V2M (bottom, blue). Presynaptic spikes above uIPSCs in black. Grey traces are individual sweeps. **B:** Population data of uIPSC amplitude (top), failure rate and CV (bottom) recorded at the three synapses in control (open symbols) and in the presence of AM-251 (filled symbols). Color code as in A. **C:** Schematic summary of the main finding of this study. Grey area represents the specific expression of tonic CB1 signaling conferring weaker inhibition from CB1 BCs onto PNs in L2/3 of V2M.

Altogether, these results indicate that tonic CB1 signaling is responsible for decreasing the strength of GABAergic synapses made by CB1-expressing interneurons selectively in L2/3 of the associative cortex V2M. Strikingly, this mechanism controls GABA release probability differentially at synapses from the same interneuron, whose axon originates in L2/3 but target different postsynaptic PNs in either L2/3 or L4.

### Spontaneous activity of PNs is higher in V2M as compared to V1

Our previous results indicate that CB1 BCs operate a differential control of PN perisomatic region in V2M as compared to V1 through tonic CB1 signaling. Tonic inhibition of GABA release from CB1 BCs should in principle result in higher output activity of PNs. To test this hypothesis, we performed *in vivo* 2-photon (2P) Ca^2+^ imaging in awake mice, free to locomote on a circular treadmill (Figure 5A). We measured spontaneous PN activity in L2/3 in either V1 or V2M in CB1 td-Tomato mice. We injected either V1 or V2M with an adeno-associated viral (AAV) vector expressing the genetically encoded Ca^2+^ indicator GCaMP6f under the pan-neuronal promoter synapsin I (*SynI*). This allowed us to measure Ca^2+^ transients as a proxy for neuronal firing from putative PNs and td-Tomato-expressing CB1 BCs (Figure 5B). We found that the overall mean change in fluorescence (ΔF/F_0_) was much higher in V2M than in V1 (mean ΔF/F_0_ = 0.17 ± 0.02 vs. 0.31 ± 0.012, V1 vs. V2M, respectively; n = 7 and 6 mice in V1 and V2M, respectively; p= 0.0023; Mann–Whitney U test, Figure 5C-D) suggesting differences in spontaneous activity of PNs across the two brain regions. We found no differences in mean movement percentage (21.70 ± 3.461% vs 23.75 ± 0.9961% in V1 and V2M, respectively; p= 0.2949; Mann– Whitney U test) and speed (0.4391 ± 0.05976 cm/s vs 0.5161 ± 0.09860 cm/s in V1 and V2 M, respectively; p= 0.6282; Mann–Whitney U test) between the two animal groups. This indicates that the differences in animal state are likely not the underlying cause of increased neuronal activity in V2M. We then restricted our analysis on td-Tomato-expressing neurons, and we found that CB1 INs did not exhibit statistically different activity in the two visual areas (mean ΔF/F_0_ = 0.27 ± 0.04 vs. 0.38 ± 0.04, V1 vs. V2M, respectively; n = 7 and 6 mice in V1 and V2M, respectively; p= 0.10; Mann–Whitney U test, Figure 5E-F). This suggests that increased activity in V2M was overall restricted to PNs.

**Figure 5.**
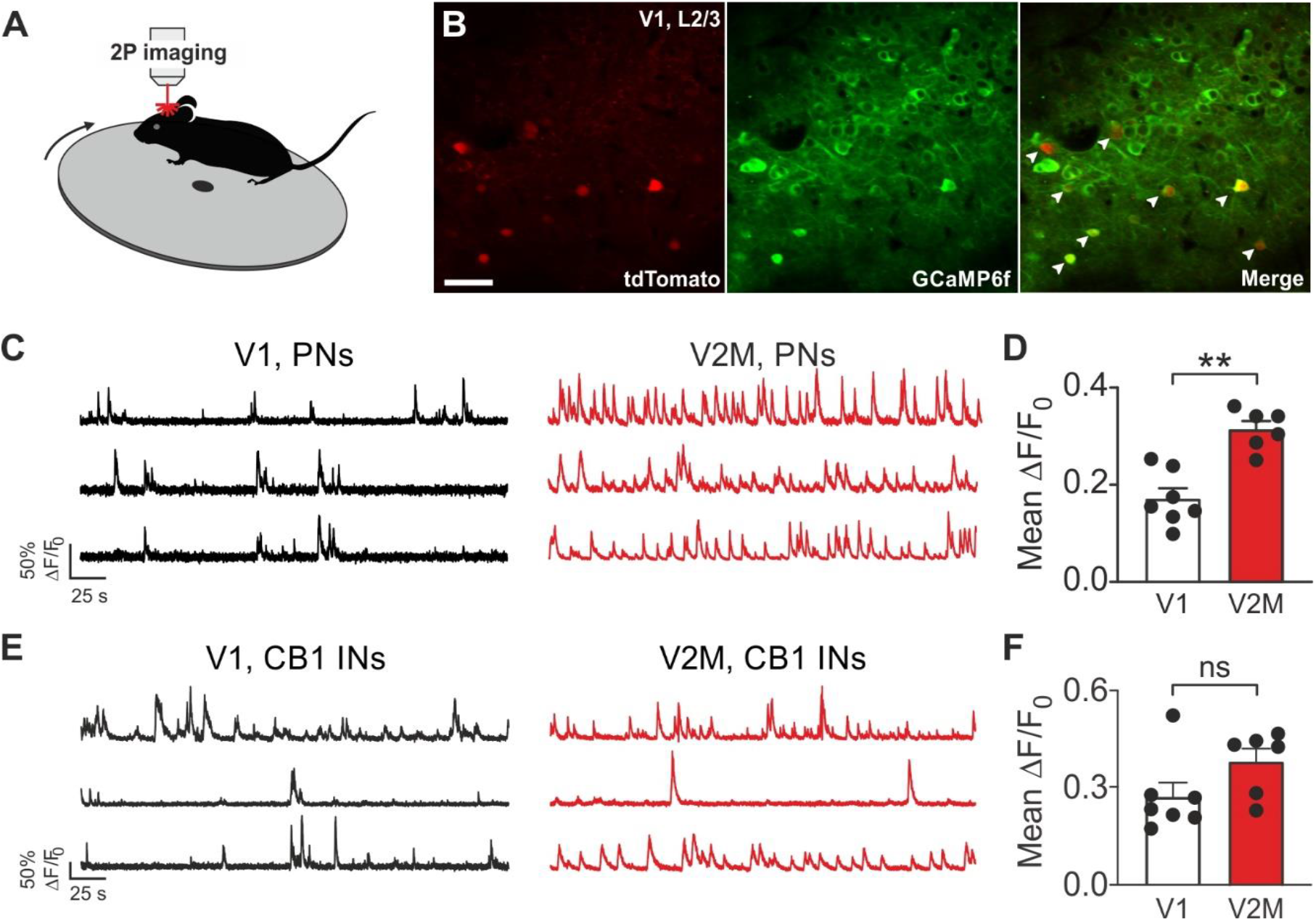
Spontaneous activity of PNs is higher in V2M as compared to V1. **A:** Schematic of 2P imaging recordings in awake mice free to locomote on a rotating disk. **B:** Representative, average intensity projection images obtained in L2/3 of V1 from a CB1-tdTomato mouse. Red indicates tdTomato expression (left), green shows GCaMP6f expression (middle). The right panel illustrates the overlay of the red and green channels. Arrowheads point to CB1-expressing neurons. Scale bar: 50 µm. **C:** Representative 2P Ca^2+^ fluorescence traces from three PNs in V1 (black, left) and in V2M (red, right). **D:** Population histogram of the mean ΔF/F_0_ in V1 (white column) and V2M (red column). Each dot represents an individual mouse. **E-F:** Same as in C-D but on tdTomato expressing interneurons.

Altogether, these results indicate that PNs in V2M fire more than in V1 in basal conditions, consistent with tonic CB1 signaling at a prominent GABAergic synapse impinging their perisomatic region.

### Visual area-specific tonic CB1 signaling underlies higher activity in V2M than V1

Higher activity of V2M PNs than V1 PNS could derive from a differential modulation of perisomatic inhibitory control. We have shown that V2M L2/3 PNs were targeted by weaker GABAergic synapses because of persistent CB1 signaling. To confirm that higher PN activity *in vivo* was due to visual area-specific tonic CB1 signaling, we injected mice with the specific CB1 inverse agonist SR 141716A (Rimonabant; 5 mg/kg intraperitoneal) (Saravia et al., 2017). Mice were injected with either vehicle or Rimonabant (see STAR Methods) and imaging sessions started 30 minutes up to 1 hour post-injection. We found that Rimonabant did not alter L2/3 neuronal spontaneous activity in V1 (mean ΔF/F_0_ = 0.1904 ± 0.03102 vs. 0.1954 ± 0.05205, V1, vehicle vs. V1, Rimonabant, respectively; n = 6 and 5 mice in V1, vehicle vs. V1, Rimonabant, respectively; p>0.9999; Wilcoxon matched-pairs signed rank test, Figure 6A-B). In support of tonic CB1 signaling, the activity of V2M PNs was reduced by ∼40% by Rimonabant (mean ΔF/F_0_ = 0.39 ± 0.06 vs. 0.23 ± 0.04, V2, vehicle vs. V2, Rimonabant, respectively; n = 6 and 6 mice in V1, vehicle vs. V1, Rimonabant, respectively; p= 0.0312; Wilcoxon matched-pairs signed rank test, Figure 6A-B). These results indicate that the overall higher activity of L2/3 PNs in V2M was due to CB1 signaling.

**Figure 6.**
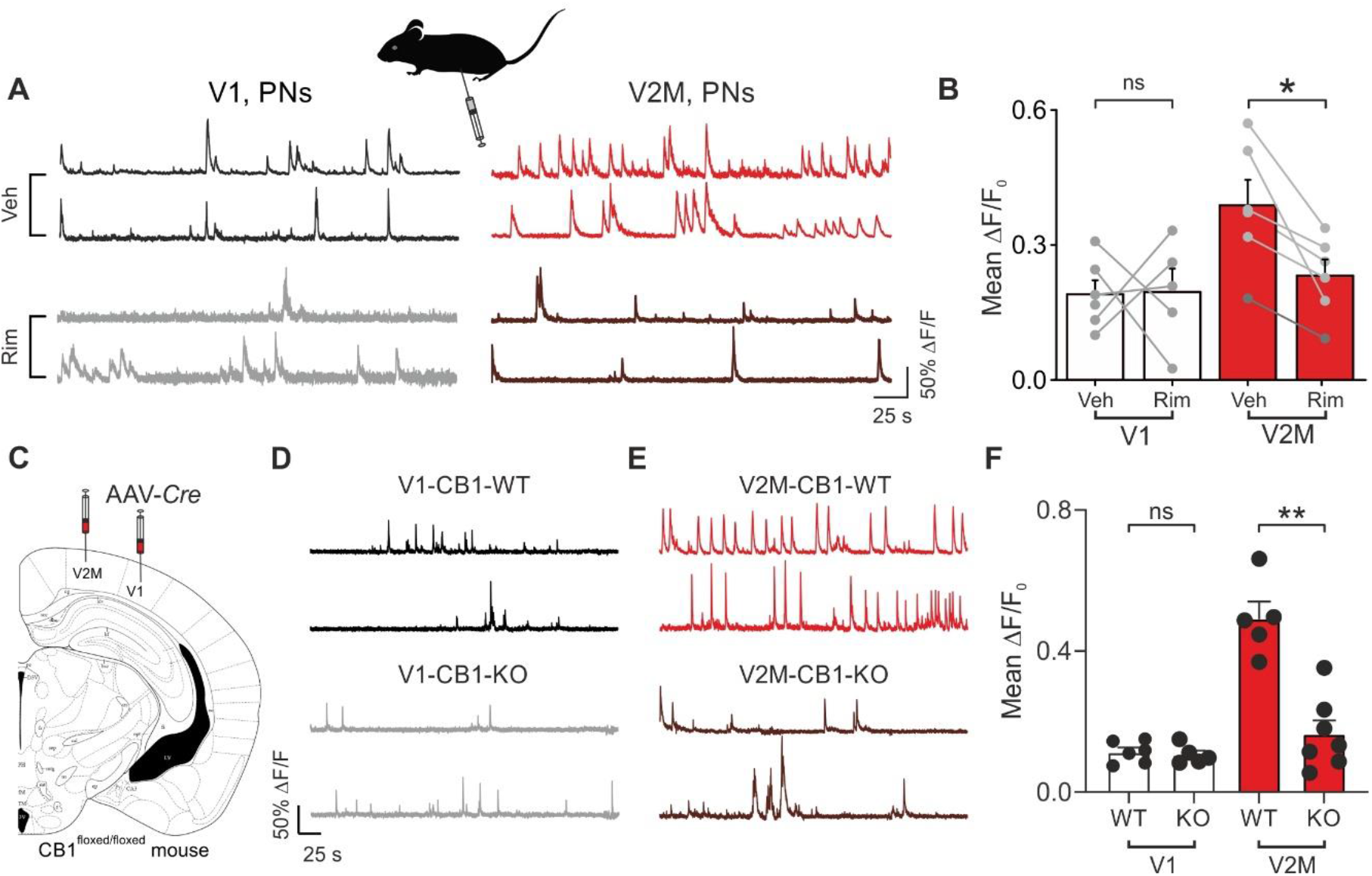
Visual area-specific tonic CB1 signaling underlies higher activity in V2M than V1. **A, Top:** Representative 2P Ca^2+^ fluorescence traces from two PNs in V1 (black, left) and in V2M (red, right) in two control mice (one for each visual area), injected with vehicle solution. **A, Bottom:** Representative 2P Ca^2+^ fluorescence traces from two PNs in V1 (gray, left) and in V2M (brown, right) two mice (one for each visual area), injected with the CB1 antagonist Rimonabant. **B:** Population histogram of the mean ΔF/F_0_ in V1 (white columns) and V2M (red columns) for vehicle- and Rimonabant-injected mice. Each grey dot represents an individual mouse. All mice were CB1- tdTomato mice, except the one represented by a darker dot, which refers to a CB1^fl/fl^ mouse. **C:** Schematic of the local genetic CB1 deletion in the two visual areas. Adult CB1^fl/fl^ mice were stereotaxically injected in either V1 or V2M with AAV-*Cre* or control (AAV-tdTomato) vectors. **D:** Representative 2P Ca^2+^ fluorescence traces from two PNs in V1 injected with the control AAV (V1-CB1- WT, top) and AAV-*Cre* (V1-CB1-KO) vectors (bottom). **E:** Same as in D, but for two mice injected in V2M. **F:** Population graph illustrating the mean ΔF/F_0_ in V1 (white columns) and V2M (red columns) for mice (black dots) injected with control or *Cre* vectors.

Systemic pharmacological blockade of CB1 does not exclude that the reduction of V2M neuronal activity was due to a global CB1 effect. In order to downregulate CB1 function in adult mice locally in either V1 or V2M only, we injected AAVs expressing the recombinase *Cre* under the pCAG promoter in mice carrying a loxP-flanked CB1 gene (CB1-flox) in either cortical area *CB1*^floxed/floxed^ (Figure 6C). This strategy was shown to knock-down successfully CB1 expression in the targeted areas (Soria-Gomez et al., 2014;Soria-Gomez et al., 2015). In fact, five weeks post-virus injection resulted in a significant reduction of CB1 immunoreactivity in either V1 or V2M (p= 0.0313, Wilcoxon matched-pairs signed rank test; Figure S7. We found that local genetic knockdown of CB1 did not alter L2/3 PN activity in V1 (mean ΔF/F_0_ = 0.11 ± 0.012 vs. 0.10 ± 0.012, V1-CB1_WT vs. V1-CB1-KO, respectively; n = 6 and 5 mice, p= 0.93; Mann–Whitney U test, Figure 6D-F) but strongly reduced the firing of PNs in V2M (mean ΔF/F_0_ = 0.49 ± 0.05 vs. 0.16 ± 0.038, V2M-CB1_WT vs. V2M-CB1-KO, respectively; n = 5 and 7 mice, p= 0.0025; Mann–Whitney U test, Figure 6D-F).

These results demonstrate the existence of tonic CB1 signaling *in vivo*, which is restricted to V2M. This modulation strongly affects neuronal activity specifically in this higher order visual area, likely by persistently reducing GABA release from CB1 BCs. Overall, these results indicate that these BCs operate a strong control of PN firing.

### Modulation of perisomatic inhibition from CB1 BCs affects correlated activity of PNs selectively in V2M

Perisomatic inhibition can in principle affect the degree of correlated PN activity (Pouille and Scanziani, 2001;Gabernet et al., 2005;Freund and Katona, 2007;Manseau et al., 2010;Buzsaki, 2010;Lourenco et al., 2014;Lourenco et al., 2020a). However, we found that the relative strength of perisomatic inhibition from CB1 BCs is strikingly different between V1 and V2M. Therefore, we extracted the relative timing of the recorded Ca^2+^ events between PNs in our 2P imaging experiments and quantified pairwise correlations of PNs in V1 and V2M. To do so, we applied the spike time tiling coefficient (STTC) method to our Ca^2+^ imaging data and reduce possible confounding effects linked to differences in firing rates (see STAR Methods) (Cutts and Eglen, 2014). We first deconvolved fluorescence traces to extract putative times of spike events as previously done (see STAR Methods) (Stringer and Pachitariu, 2019). We found that neurons in V2M (characterized by higher activity) exhibited less correlated activity of Ca^2+^ events as compared to V1 (STTC: 0.057 ± 0.004 vs 0.047 ± 0.004, V1 vs. V2M; n = 70 and 62 time series; p = 0.033; Mann–Whitney U test, Figure 7A-B).

**Figure 7.**
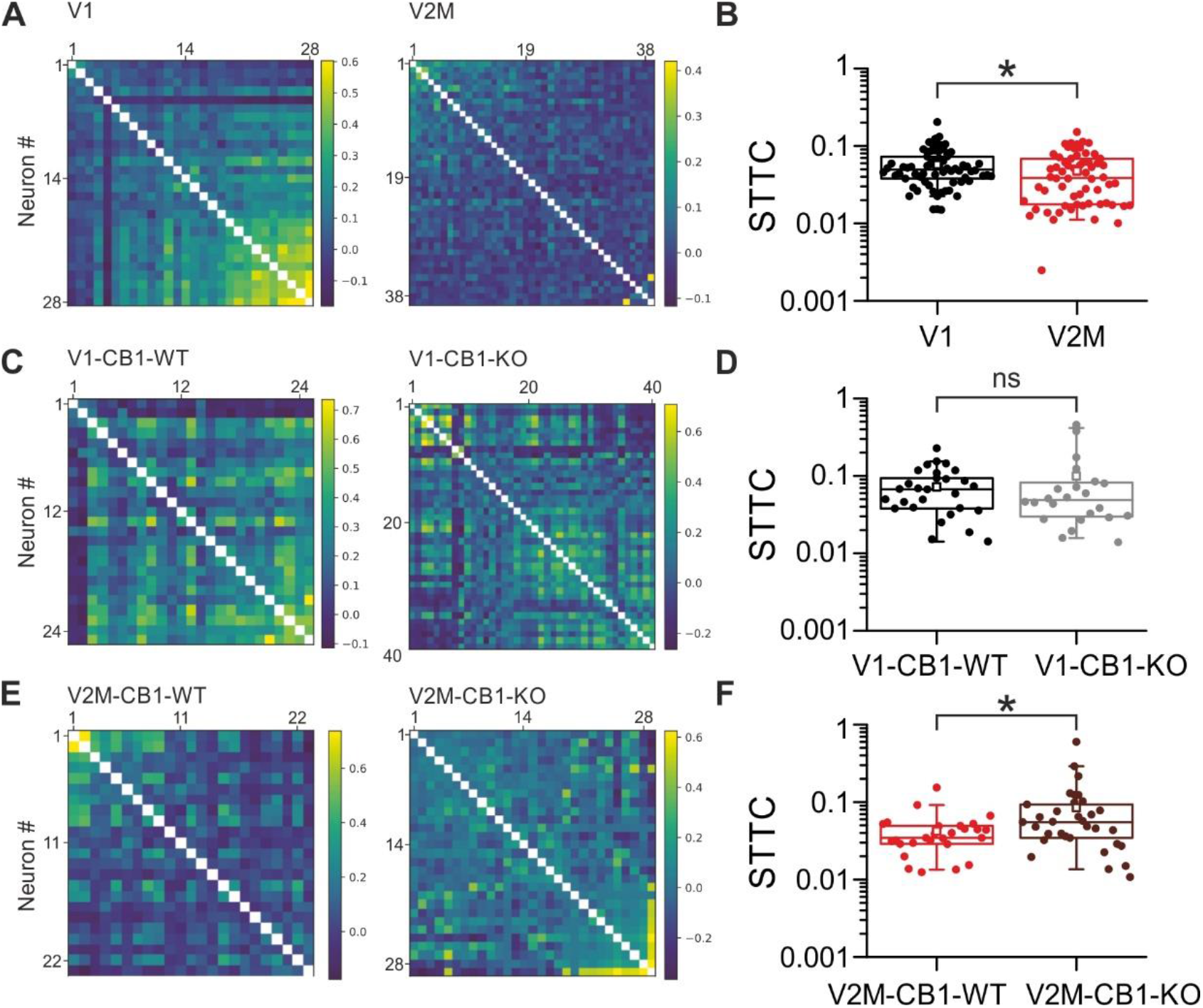
Modulation of perisomatic inhibition from CB1 BCs affects correlated activity of PNs selectively in V2M. **A:** Pairwise correlation matrices of two representative time series in V1 (left) and V2M (right) from CB1-tdTomato mice. The heatmaps indicate low (blue), intermediate (green) and high (yellow) levels of correlated activity. **B:** Population data of the spike time tiling coefficient (STTC) in V1 (black) and V2M (red). STTC is a coefficient quantifying the degree of correlated activity of 2P Ca^2+^ imaging neurons. Each dot is a time-series. **C-F:** Same as in A-B but for CB1^fl/fl^ mice injected with control AAV (CB1-WT) and AVV-Cre (CB1-KO) vectors in V1 (C-D) and V2M (E-F). Color code for different viral vectors and visual areas in D and F as in Fig. 6 D-E.

To test whether the smaller degree of correlation was due to perisomatic control by CB1 BCs, we analyzed STTC in CB1*^floxed/floxed^* mice injected with AAV-Cre vectors. We found that STTC values were unchanged in V1 of mice injected with either control or Cre*-*expressing viruses (STTC: 0.072 ± 0.01 vs 0.099 ± 0.026, V1-CB1-WT vs. V1-CB1-KO; n = 29 and 24 time series; p = 0.67; Mann–Whitney U test, Figure 7C-D). Interestingly, in addition to reducing overall PN activity (Figure 6), knocking out CB1 in V2M, resulted in a significant increase of correlated activity in this visual area (STTC: 0.042 ± 0.005 vs 0.08 ± 0.019, V2M-CB1-WT vs. V2M-CB1-KO; n = 27 and 33 time series; p = 0.023; Mann–Whitney U test, Figure 7E-F).

In sum, these data indicate that CB1 BCs contribute significantly to orchestrate cortical networks. More strikingly, these results indicate that tonic CB1 signaling represents a simple, albeit powerful mechanism to controls the level of activity and coordination of PNs in different cortical areas.

## Discussion

In this study, we examined the functional connectivity pattern of CB1 BCs in the mouse visual cortex in slices and *in vivo*. CB1 BCs are elusive elements of the cortical microcircuit. Their functional properties are much less known as compared to other inhibitory cell types, such as dendritic targeting somatostatin (SST) interneurons and PV cells. We found that CB1 expression was higher in V2M across L2-5 as compared to V1. Moreover, we found that CB1-expressing BCs exhibited similar electrophysiological and synaptic properties in the two visual cortical areas. However, surprisingly, they possessed different anatomical and connectivity patterns in V1 vs. V2M. Overall, CB1 BCs in L2/3 of V2M had a much lower efficacy of synaptic release, due to a persistently active CB1 signaling in this specific layer and cortical visual area (Figure 4C). This area-specific connectivity and eCB modulation was responsible of a higher although less coordinated activity of PNs in V2M, as compared to V1.

Our anatomical data are in good agreement with previously reported differences of expression of CB1 in different cortical areas (Yoneda et al., 2013). More CB1 in V2M could be due to: *i)* differences in the number of CB1 interneurons; *ii)* differences in their projections; *iii)* increased expression of CB1 in each axonal terminal. We found support for both i) and ii): we report a significant larger proportion of CB1 interneurons, and high levels of axonal innervation originating in L2/3 and invading L4, only in V2M. Regarding iii), our results cannot exclude higher expression of CB1 in V2M, and future pharmacological studies are required to directly address this question. However, it is notable that comparable levels of CB1 in L2/3 and L4 of V2M yielded completely different CB1- dependent modulation of synaptic transmission and plasticity. Moreover, it was shown that tonic CB1- mediated modulation of GABA release is independent of the expression levels of CB1 on axon terminals (Ladarre et al., 2014). Higher CB1 expression in V2M could serve multiple cellular functions (Ladarre et al., 2014), possibly via differential expression in specific subcellular compartments, such as mitochondria (Hebert-Chatelain et al., 2016).

The V2M-specific innervation pattern onto L4 is particularly intriguing. The lack of infra-layer innervation from CB1 BCs in V1 could be due to a consistent inability of filling axons with biocytin, or a bias in selecting CB1 BCs in specific cortical areas. This is unlikely for at least two reasons. First, the prominent V2M L4 innervation in filled neurons is in agreement with the CB1 staining pattern. Second, the probability of finding infra-laminar connected pairs is dramatically different in the two visual areas; we could find very few connected pairs when the postsynaptic PN was in L4 of V1 (5 pairs out of 74 double recordings). Here we provide evidence of a specific connectivity logic of CB1 BCs from L2/3 in distinct visual areas. The specific infra-laminar projection of CB1 BCs in V2M might reveal different circuit motifs in different cortical areas. Indeed, L4 activation by thalamo-cortical fibers is relayed to L2/3 where it could generate a feedback inhibitory loop operated by CB1 BCs only in associative cortices. This associative area-specific routing of intracortical information could have important consequences in sensory perception.

Differences in anatomical parameters, such as dendritic and axonal projections have traditionally been used to distinguish different interneuron subclasses (Markram et al., 2004;Ascoli et al., 2008;Naka et al., 2019). Accordingly, different morphologically identified cell types exhibit specific electrophysiological properties (Markram et al., 2004). Here we found that, despite the different axonal projections in the two visual areas, CB1 BCs exhibited a similar electrophysiological signature. Moreover, GABAergic synaptic transmission from CB1 BCs share many similarities in the two visual areas, such as high failure rate and variable short-term dynamics, consistent with their counterparts in the hippocampus (Hefft and Jonas, 2005;Neu et al., 2007;Daw et al., 2009), amygdala (Rovira-Esteban et al., 2017) and other cortical areas (Galarreta et al., 2008). This argues against the existence of distinct subtypes of cortical CB1 BCs in different visual areas, although a thorough molecular investigation using single-cell transcriptomics will be important to pinpoint possible selective expression profiles. Importantly, however, GABAergic transmission was much weaker onto L2/3 PNs of V2M, but this layer- and visual area-specific synaptic efficacy was erased by blocking CB1. This indicates that the different inhibitory strength exhibited in L2/3 of V2M did not depend on the specificity of pre- and postsynaptic molecular architecture (Eggermann et al., 2011), but it was due to persistent modulation of GABA release from CB1 BCs, only in this layer of this associative visual area, resulting in functional synaptic diversity and target specificity.

Despite an increase in uIPSC amplitude and a decrease in failure-rate, the lack of change in PPR in the presence of AM251 is unexpected, but also consistent with other studies (Kim and Alger, 2001;Rovira-Esteban et al., 2017). Thus, PPR with two consecutive IPSCs may not represent an appropriate measure of presynaptic release probability at these synapses. Indeed, using PPR to infer presynaptic release probability of GABAergic synapses has been questioned in the hippocampus, due to spurious facilitation caused by occasional synaptic failure (Kim and Alger, 2001). Importantly, however, AM-251 application decreased uIPSC failure-rate, consistent with a presynaptic modulation by CB1 (Neu et al., 2007).

Despite selective tonic CB1 modulation, CB1-dependent DSI magnitude was similar at the three tested connections. This indicates that tonic CB1 signaling in V2M does not saturate the receptor, which is still sensitive to depolarization-induced, on-demand production of eCBs. Importantly, however, despite the *relative* DSI amplitude was similar in V1 and V2M, it is noteworthy to stress here that, from the perspective of single PNs, DSI produced a massive reduction of PN perisomatic inhibition in L2/3 of V1 (synaptic currents went from several hundred pA to zero). In contrast, L2/3 PNs of V2M sensed a reduction of perisomatic inhibition from CB1 INs, which was already weak (∼6 fold less powerful) already before DSI stimuli. This differential absolute change of acute eCB modulation of inhibitory transmission from CB1 interneurons will likely produce distinct effects in the output spiking properties of single PNs in the two cortical areas.

Importantly, CB1-dependent tonic reduction of inhibition has been reported at GABAergic synapses in the hippocampus (Losonczy et al., 2004;Foldy et al., 2006;Neu et al., 2007), although the existence of tonic CB1 signaling was disputed (Castillo et al., 2012). This has raised the possibility that tonic inhibition could depend on the health of the slice preparation and/or recording conditions. Yet, here we demonstrate in adult tissue that the same CB1 neuron could be responsible for phasic and tonic modulation at different synapses. Moreover, we found that tonic CB1 modulation had a strong impact in modulating *in vivo* PN firing behavior in V2M selectively.

Tonic CB1 activation could be due to different mechanisms, including a constitutively active receptor in the absence of a natural ligand (Leterrier et al., 2004;Losonczy et al., 2004), or a persistently activated receptor by a tonic presence and/or synthesis of eCBs (Neu et al., 2007). It has been suggested that this tonic eCB mobilization from postsynaptic PNs is regulated by mGluRs and muscarinic receptors (Foldy et al., 2007). Yet, we believe that this tonic modulation of CB1 is due to a constant presence of eCBs. Indeed, within the same cortical area (V2M) the same axons exhibited strong and absent tonic CB1 signaling, in L2/3 and L4, respectively, arguing against a target-specific endogenous state of CB1. It will be interesting to determine the possible sources of eCBs: these could be postsynaptic neurons or non-neuronal elements in the neuropil (such as astrocytes). Alternatively, tonic CB1 activation could derive from a layer-specific absence of enzymatic eCB degradation (Ladarre et al., 2014).

Independently of the actual underlying molecular mechanism, it has been shown that tonic CB1 activity can be overrun by high frequency firing of the presynaptic interneurons (Chen et al., 2003;Chen et al., 2007;Foldy et al., 2007) or could be facilitated by presynaptic activity (Zhu and Lovinger, 2007;Heifets et al., 2008;Lourenco et al., 2010). Here we show that the strength of CB1 in- mediated inhibition depends on the activity state of these inhibitory neurons. Maturation of the visual cortex and stress conditions could alter CB1 expression and thus represent factors altering CB1 signaling (Jiang et al., 2010;Wamsteeker Cusulin et al., 2014).

Weak perisomatic inhibition in V2M might be used as a strategy to modulate postsynaptic PN firing. Indeed, we observed a much higher spontaneous *in vivo* activity of PNs in this associative area, as opposed to V1. This is consistent with overall decreased inhibition onto PNs. In sensory cortices, L2/3 PNs exhibit low-frequency activity, suggesting that they integrate sensory input using sparse coding (Petersen and Crochet, 2013). We found a stark difference of activity of L2/3 PNs in adjacent visual cortical areas. One can thus speculate that higher-order visual cortical regions encode sensory information using a different computational strategy. Future experiments will define the actual role of this difference in firing in the hierarchical flow of sensory information involving primary and associative cortical areas (Glickfeld and Olsen, 2017;Minderer et al., 2019;Jin and Glickfeld, 2020;Siegle et al., 2021).

Yet, is this visual area-specific PN activity level dependent on the strength of GABAergic neurotransmission onto PNs? In particular, do CB1 BCs contribute to set the activity level of PNs in different cortical areas? We found that pharmacological blockade and genetic deletion of CB1 reduced the activity of PNs in V2M to levels similar to V1. The decrease of neuronal activity observed in V2M after pharmacological CB1 blockade could be ascribed to a brain-wide -or even peripheral- effect. However, the virtually identical results were obtained by acute and localized genetic deletion of CB1 in V1 or V2M in adult mice, thus strengthening our interpretation that this difference in neuronal activity is due to tonic CB1 signaling. This result strongly suggests that CB1 INs control the output activity of PNs. Moreover, this result reveals the presence of tonic CB1 signaling in a specific cortical area (V2M), in good agreement with our anatomical and synaptic findings (Figs. 1-4).

The difference in PN firing in V1 and V2M was not associated with significant firing alterations of CB1 INs. This suggests that these cells are likely not recruited by local PNs, which exhibit higher activity in the visual areas. Moreover, the lack of difference in firing frequency between CB1 INs in V1 and V2M suggests a functional decoupling between their firing behavior and synaptic release of GABA. Indeed, the overall spontaneous activity of CB1 INs may not be sufficient to overcome tonic CB1 inhibition of GABA release.

One hallmark of perisomatic inhibition is the ability of controlling the timing of output spiking of PNs. Indeed, reliable and strong perisomatic inhibition from PV BCs was shown to reduce the jitter and increase the reliability of their postsynaptic targets (Pouille and Scanziani, 2001;Gabernet et al., 2005;Lourenco et al., 2014;Lourenco et al., 2020a), thus ending up synchronizing large populations of PNs and favoring the emergence of rhythmic correlated activity (Bartos et al., 2007;Buzsaki and Silva, 2012). The significant reduction in pairwise correlation that we observed in V2M is consistent with decreased strength of perisomatic inhibition. Indeed, correlated activity is related to higher firing frequency (de la Rocha et al., 2007), but we observed a lower synchrony in V2M in the presence of higher firing activity. Local genetic deletion of CB1 did not affect synchrony in V1 but it increased coordinated activity in V2M. Altogether, these results are consistent with a significant role played by CB1 BCs in orchestrating cortical networks. This is intriguing, given the paucity of this cell type and their loose-coupled synapse as compared to the more pervasive PV cells, characterized also by reliable and fast synaptic transmission (Hefft and Jonas, 2005;Freund and Katona, 2007;Deleuze et al., 2019). Future experiments will be necessary to unravel the actual synchronization role played by CB1 BCs with a better temporal resolution, to reveal whether these BCs are involved in fast network oscillations. The highly specific inhibitory modulation of PN output can profoundly affect the participation and orchestration of populations of PNs to relevant network oscillations, proposed to underlie several cognitive functions, including sensory perceptions (Buzsaki, 2010;Siegle et al., 2014). Interestingly, it has been recently shown that hippocampal CCK/CB1 and PV BCs play a complementary role during fast network activity, due to mutual inhibitory connections between these IN subclasses (Dudok et al., 2021). Whether this scheme applies also to different visual cortical areas is not known. Here, it is tempting to speculate that given our effect of CB1 tonic modulation selectively in V2M, perisomatic inhibition from CB1 BCs play a predominant role in this specific cortical area.

In sum, here we found that different morpho-functional and connectivity properties of a specific GABAergic IN subtype governs the activity level of a visual cortical area, suggesting that distinct circuit blueprints can define the function of specific cortical areas during sensory perception.

## Experimental Procedures

### Animals

Experimental procedures followed French and European guidelines for animal experimentation and in compliance with the institutional animal welfare guidelines of the Paris Brain Institute. Experiments for paired recordings were performed on both sexes aged between P30 and P40 CB1-tdTomato mice (Winters et al., 2012). In some experiments, CB1-tdTomato mice were crossed with GAD67-GFP mice to identify CB1-expressing INs for cell counting. In experiments for Figure 1, C57BL/6J wild-type were purchased from Janvier laboratories. Mice were housed in an animal facility with a 12h light/dark cycle, with food and water available *ad libitum*. Viral injections and chronic cranial windows for 2P Ca^2+^ imaging experiments *in vivo* were performed on two months old male mice of CB1-tdTomato or CB1^floxed/floxed^ mice and imaging sessions started following habituation, four to five weeks post-surgery.

### Immunohistochemistry and cell counting

All animals were deeply anesthetized with ketamine-xylazine and transcardially perfused first with cold PBS (20ml) followed by 30-40 ml of cold 4% PFA (paraformaldehyde, diluted in PBS). Brains were removed and post-fixed in 4% PFA overnight at 4°C. For cryoprotection they were next put in 30% sucrose (diluted in PBS) overnight and frozen at - 45°C in isopentane. The brains were cut with a freezing microtome (Thermo Fisher), with nominal section thickness set to 20 µm. After rinsing with PBS, slices where incubated 2h at room temperature in 0.3% PBT (0.3% Triton X in PBS) and 10% BSA blocking solution. Primary antibodies diluted in 0.3% PBT and 0.1% NGS (normal goat serum) were incubated overnight at 4°C. The following antibodies were used: anti-CB1 (Frontiers institute, goat 1:400), anti-GFP (Millipore, MAB 3580 mouse 1:500), anti-DsRed (Clontech, rabbit 1:500), anti-PV (Sigma PARV-19, mouse 1:1000) and anti-SST (Santa Cruz G10 sc-55565, mouse 1:250). The CB1 antibody was raised against a 31 amino acid C-terminal sequence of the receptor, and has been shown to mainly stain GABAergic axonal fibres, resulting in an immunopositive mesh (Bellochio *et al.,* 2010). Anti-GFP and anti-DsRed (i.e. tdTomato) were used to enhance the signals of GFP and tdTomato allowing us to increase the signal to noise ratio, particularly for counting somata. Slices were then rinsed with PBS and incubated for 2h at room temperature with the secondary antibodies Alexa 488 anti-goat, Alexa 488 anti-mouse and Alexa 633 anti-rabbit, all obtained from Life Technologies and diluted 1:500 in PBT 0.3%. Slices were mounted with DAPI Fluoromount (Sigma) and stocked at 4°C. Whole brain slices were imaged using an epifluorescence slice scanner (Axio scan Z1 Zeiss, magnification 20x).

### CB1 immunofluorescence pattern

Only slices in which both V1 and V2M areas were present were analyzed in order to be able to quantitatively compare the fluorescence patterns. To obtain the pattern, the “straight” option in FIJI (NIH) with a line width of 390 µm was used to calculate the grey value intensity of CB1 immuno- staining from pia to white matter. For each slice, the maximum fluorescence intensity over a length of 10 µm of either V1 and V2 was used to normalize fluorescence in the rest of the analyzed areas. Moreover, due to differences in cortical length between animals and slices, we determined cortical layer thickness by using a ratio obtained from measuring each layer in the Allen atlas with total cortical length set as 1.

### Combined Fluorescent in situ hybridization (FISH)/ Immunohistochemistry (IHC) on free-floating frozen sections

CB1 FISH/tdTomato immunofluorescence experiments were carried out as previously described (Marsicano and Lutz, 1999;Oliveira da Cruz et al., 2020). Briefly, free-floating frozen coronal sections were cut out with a cryostat (20 µm, cryostat Leica CM1950), collected in an antifreeze solution and conserved at -20°C. After inactivation of endogenous peroxidases and blocking with 3% H_2_O_2,_ the tissue was treated with Avidin/Biotin Blocking Kit (SP-2002 Vector Labs, USA) followed by an incubation overnight at 4°C with antiDsRed rabbit polyclonal primary antibody (1:1000, 632496 Takara Bio) diluted in a triton buffer (0.3% Triton X-100 diluted in PBS-DEPC). The following day, after several washes, the sections were incubated with a secondary antibody goat anti-rabbit conjugated to a horseradish peroxidase (HRP) (1:500, 7074S Cell Signaling Technology) during 2 hours at RT followed by a 10 minutes incubation at RT with TSA plus Biotin System (Biotin TSA 1:250, NEL749A001KT PerkinElmer). After several washes, the slices were fixed with 4% of formaldehyde (HT501128-4L Sigma Aldrich) for 10 minutes, incubated with 3% of H_2_O_2_ for 30 minutes at RT and blocked with 0.2M HCl for 20 minutes at RT. Then, the tissue was acetylated in 0.1 M Triethanolamine, 0.25% Acetic Anhydride for 10 minutes. Sections were hybridized overnight at 70°C with Digoxigenin (DIG)-labeled riboprobe against mouse CB1 receptor (1:1000, prepared as described in Marsicano and Lutz, 1999). After hybridization, the slices were washed with different stringency wash buffers at 70°C. Then, incubated with 3% of H_2_O_2_ for 30 minutes at RT and blocked 1 hour RT with NEN blocking buffer prepared according to the manufacturer’s protocol (FP1012 PerkinElmer). AntiDIG antibody conjugated to HRP (1:1500, 11207733910 Roche) was applied for 2 hours at RT. The signal of CB1 receptor hybridization was revealed by a TSA reaction using cyanine 3 (Cy3)-labeled tyramide (1:100 for 10 minutes, NEL744001KT PerkinElmer). After several washes, the free-floating sections were incubated 30 minutes at RT neutravidin dyligth 633 (1:500, 22844 invitrogen). Finally, the slices were incubated with DAPI (1:20000; 11530306 Fisher Scientific) diluted in PBS, following by several washes, to finally be mounted, cover slipped and imaged using an epifluorescence slice scanner (Axio scan Z1 Zeiss, magnification 20x) (see below section on “Counting CB1 INs” for analysis).

CCK FISH/tdTomato immunofluorescence experiments were carried out as previously described (Oliveira da Cruz et al., 2020). Briefly, free-floating frozen coronal sections were cut out with a cryostat (20 µm, cryostat CM1950 Leica), collected in an antifreeze solution and conserved at -20°C. After inactivation of endogenous peroxidases and blocking with 3% H_2_O_2_ the tissue was treated with Avidin/Biotin Blocking Kit (SP-2002 Vector Labs, USA) followed by an incubation overnight at 4°C with antiDsRed rabbit polyclonal primary antibody (1:1000, 632496 Takara Bio) diluted in a triton buffer (0.3% Triton X-100 diluted in PBS-DEPC). The following day, after several washes, the sections were incubated with a secondary antibody goat anti-rabbit conjugated to a horseradish peroxidase (HRP) (1:500, 7074S Cell Signaling Technology) during 2 hours at RT followed by a 10 minutes incubation at RT with TSA Biotin System (Biotin TSA 1:100, NEL700A001KT PerkinElmer). After several washes, the slices were fixed with 4% of formaldehyde (HT501128-4L Sigma Aldrich) for 10 minutes and blocked with 0.2M HCl for 20 minutes at RT. Then, the tissue was acetylated in 0.1 M Triethanolamine, 0.25%

Acetic Anhydride for 10 minutes. Sections were hybridized overnight at 60°C with Digoxigenin (DIG)- labeled riboprobe against CCK (1:1000, (Marsicano and Lutz, 1999)). After hybridization, the slices were washed with different stringency wash buffers at 65°C. Then, incubated with 3% of H_2_O_2_ for 30 minutes at RT and blocked 1 hour RT with NEN blocking buffer prepared according to the manufacturer’s protocol (FP1012 PerkinElmer). AntiDIG antibody conjugated to HRP (1:2000, 11207733910 Roche) was applied for 2 hours at RT. The signal of CCK hybridization was revealed by a TSA reaction using fluorescein isothiocyanate (FITC)-labeled tyramide (1:100 for 10 minutes, NEL741001KT Perkin Elmer). After several washes, the free-floating sections were incubated overnight at 4°C with Streptavidin-Texas Red (1:400, NEL721001EA PerkinElmer). Finally, the slices were incubated with DAPI (1:20000; 11530306 Fisher Scientific) diluted in PBS, following by several washes, to finally be mounted, coverslipped and imaged using an epifluorescence slice scanner (Axio scan Z1 Zeiss, magnification 20x) (see below section on “Counting CB1 INs” for analysis).

### Counting CB1 INs

CB1-tdTomato/GAD67-GFP positive cells were counted in both V1 and V2; for each area a region of interest (ROI) was drawn. The ITCN (Image-based Tool for Counting Nuclei) plugin in ImageJ was modified (courtesy of Brahim Abbes, Neurostack) in order to be able to count the cells and their distance from the pia simultaneously. The pia was traced on each image, and cells expressing both tdTomato and GFP were manually marked. Indeed, although the ITCN was designed to automatically count cells, the widespread expression of tdTomato in both soma and neurites made automatic detection too unreliable. The software then calculated the minimal distance from marked cells to the pia. Once these distances were established, we were able to bin cell counts in the different layers of the cortex. Layer (bin) size was determined using the online mouse reference atlas, determining the % of cortical thickness each layer represented. We then used these ratios to calculate actual layer thickness on our slices. Also, ROIs could differ widely in width, hence to compensate for this we divided the obtained laminar density by the width of the ROI.

For the CB1-tdTomato/5HT3-GFP, CB1-TdTomato/Cnr1 and CB1-tdTomato/CCK positive cells count, the same plugin was used with a new feature added (Neurostack, courtesy of Brahim Abbes): CB1- tdTomato cell area. Layer size and thickness were then determined in the same way as above.

### In Vitro Slice Preparation for Electrophysiological Recordings

Coronal slices (350 µm thick) of visual cortex were obtained from mice of both sexes aged between P30 and P40. The area of interest was identified using the Allen adult mouse brain reference atlas. Animals were deeply anaesthetized with a mix containing 120 mg/kg ketamine and 24 Xylazine mg/kg of body weight (in 0.9% NaCl). A transcardiac perfusion was performed using an ice cold “cutting” solution containing the following (in mM): 126 choline chloride, 16 glucose, 26 NaHCO_2_, 2.5 KCl, 1.25 NaH2PO_4_, 7 MgSO_4_, 0.5 CaCl_2_, (equilibrated with 95% O_2_ / 5% CO_2_). Following decapitation, brains were quickly removed and sliced with a vibratome (Leica) while immersed in ice cold cutting solution. Slices were then incubated in oxygenated (95% O_2_ / 5% CO_2_) artificial cerebrospinal fluid (ASCF) containing the following (in mM): 126 NaCl, 20 glucose, 26 NaHCO_3_, 2.5 KCl, 1.25 NaH_2_PO_4_, 1 MgSO_4_, 2CaCl_2_ (pH 7.4, 310-320mOsm/L), at 34°C for 20 min, and subsequently at room temperature before transferring to the recording chamber. The recording chamber was constantly perfused with warm (32°C), oxygenated ACSF at 2.5-3 mL/min.

### Electrophysiology

Synaptic currents were recorded in whole-cell voltage or current clamp mode in principal cells of either L2/3 or L4 of primary and secondary visual cortex. Excitatory cells of L2/3 were visually identified by their triangular soma and apical dendrites projecting towards the pia, while in L4 they were identified by their round soma and by verifying their regular firing properties. Meanwhile, CB1 BCs were targeted using CB1-tdTomato fluorescence elicited by a green (λ=530 nm) LED (Cairn research) coupled to the epifluorescence path of the microscope, alongside their characteristic large soma and bi- or multipolar dendritic morphology. To study the passive properties of CB1 BCs, action potential waveform and firing dynamics, electrodes were filled with an intracellular solution containing (in mM): 135 K-gluconate, 5 KCl,10 Hepes, 0.01 EGTA, 4 Mg-ATP, 0.3 Na-GTP and 10 phosphocreatine di(tris). The pH adjusted with KOH to 7.2 resulting in an osmolarity of 290-300 mOsm. Based on the Nernst equation, the estimated reversal potential for chloride (ECl) was approximately –84 mV. For these experiments, the following drugs were also present in the superfusate (in µM): 10 DNQX (Tocris), 10 gabazine, and 50 D-APV (all from Tocris).

To record GABAergic uIPSCs from paired recordings, we used a “high chloride” intracellular solution containing (in mM): 70 K-gluconate, 70 KCl, 10 Hepes, 0.2 EGTA, 2 MgCl_2_, 0.5 CaCl_2_, 4 Mg-ATP, 0.3 Na-GTP, 5 phosphocreatine di(tris); again, the pH was adjusted to 7.2 with KOH and resulted in an osmolarity of 290-300 mOsm. For this solution, the ECl was calculated to be at ∼-13 mV based on the Nernst equation, which means that when clamping the cell at -70mV, activation of GABAA receptors resulted in inward currents. We confirmed that the currents were GABAergic by demonstrating they were unaffected by DNQX (10 µM) (Tocris Bioscience) and extinguished by gabazine (10 µM; *data not shown*). In most paired-recording experiments, the ACSF was left drug-free as CB1 BCs were reliably targeted. Signals were amplified using a Multiclamp 700B patch-clamp amplifier (Axon Instruments), digitized with a Digidata 1440A (Axon Instruments), sampled at 50 kHz and filtered at 2 kHz or 10 kHz, respectively for voltage and current clamp recordings.

### Stimulation protocols

pClamp v. 10.3 (Axon instruments) was used to record the signal and generate stimulation protocols. All voltage-clamp protocols contained a 5 mV step used to monitor the series resistance (Rs), which was kept under 25 MΩ. Recordings in which the Rs had deviated by more than 20% were discarded. For paired recordings, a brief pulse was used to elicit a single action current, followed by a train of 5 pulses at 50 Hz. This pattern was repeated every 5s (0.2 Hz). In Figure 3, in order to test for presynaptic modulation, four trains of action potentials (10 pulses at 50 Hz, inter-train interval of 300ms) were elicited in the presynaptic CB1 BC.

*DSI:* DSI was induced in voltage clamp by holding the post-synaptic cell at 0 mV for 5 s in the post- synaptic cell. Because we encountered high failure rates and no clear trend in use-dependent short-term plasticity (see PPR in Figure 2), we averaged the uIPSC amplitudes of the entire train 5 pulses at 50Hz to obtain a single amplitude for each sweep, allowing us to have stable baselines and time-courses following depolarisation. Between three and five depolarisations were performed on each cell, separated by at least 2 min.

*AM251 pharmacology:* in experiments using the CB1 antagonist AM251 (3 µM) (Tocris and AdooQ biosciences), the drug (diluted in DMSO, less than 0.1% in ACSF) was added directly to the resting chamber.

### Data Analysis for Electrophysiological Recordings

*Action potential (AP) waveform and firing profile:* AP waveform was analyzed using Clampfit 10.3 (Molecular Devices). The threshold was defined as the first moment in which the slope of the action potential crosses a threshold typically set at 30 mV/ ms, then determining the rise time and peak of the AP. The adaptation coefficient was determined at max firing rate, by dividing the steady state spike instantaneous frequency (last two APs of depolarizing step) by the initial instantaneous frequency (first two APs of depolarizing step). Input resistance was measured in the late portion of the membrane potential relaxation from a step current injection of –100 pA, while the membrane time constant (tau) was obtained from fitting a single exponential to the early portion of the step until relaxation. Using these parameters, the capacitance was obtained arithmetically (C=tau/R). Vrest was measured during stable current clamp recordings without current injection. The relative variability of the inter-spike interval was determined by computing the coefficient of variation of the times between two consecutive APs. The full width at half maximum (FWHM) was determined by determining the width of the AP at half of its total amplitude (from threshold to peak).

uIPSC amplitudes were obtained using a custom script in MATLAB. Failures were defined as any value inferior to twice the standard deviation of the noise.

### Morphological reconstruction

*Biocytin Fills:* To reliably reconstruct the fine axonal branches of mature CB1 INs, dedicated experiments were performed. Biocytin (Sigma) was added to the intracellular solution at a high concentration (0.5g/100ml) (Jiang et al., 2015), which required extensive sonication. To avoid excessive degradation of fragile molecules such as ATP, sonication was performed in an ice bath. The intracellular solution was then filtered twice to prevent the presence of undissolved lumps of biocytin in the patch pipette. Extra care was applied in verifying that the micromanipulators and slice were stable for recordings of at least 1h. During that time, neurons were injected with large depolarizing currents in current clamp mode for fifteen times (100ms, 1-2nA, 1Hz). At the end of recordings, the patch pipette was removed carefully with the aim of resealing the cell properly, equivalent to obtaining an inside out patch. The slice was then left in the recording chamber for a further 5-10 min to allow further diffusion. Slices were then fixed with 4% paraformaldehyde in phosphate buffer saline (PBS, Sigma) for at least 48 h. Following fixation, slices were incubated with the avidin-biotin complex (Vector Labs) and a high concentration of detergent (Triton-X100, 5%) for at least two days before staining with 3,3′Diaminobenzidine (DAB, AbCam).

Cells were then reconstructed and cortical layers delimited using neurolucida 7 (MBF Bioscience) and the most up to date mouse atlas (Allen Institute). Because cortical layer thickness differs within and across areas, we normalized neurite lengths relative to layer thickness to obtain the most accurate measure of density in each layer using an arithmetic method (Bortone et al., 2014). To obtain heat maps, we imported reconstructions in Illustrator (Adobe) and aligned the soma horizontally, and pia and white matter vertically. From there, individual bitmaps were generated separating dendrites and axons. These were subsequently blurred in ImageJ (NIH) using a gaussian filter with a radius equivalent to 20 µm. The contrast of blurred images was then adjusted to obtain the highest possible pixel intensity, and were then overlapped and averaged. The resulting group average image was also adjusted to the highest pixel intensity, and a lookup table (ImageJ’s “Fire”, inverted) was applied to color code the density of neurites across cortical layers.

### Virus injections and chronic cranial window preparation

Virus injections and implantation of the cranial windows were performed as previously described (Koukouli et al., 2017). Mice were anesthetized using a mixture of ketamine (Imalgen 1000; Rhone Mérieux) and xylazine (Rompun; Bayer AG), 10 ml/kg i.p. and placed into a stereotaxic frame. The body temperature was maintained at ∼37 °C using a regulated heating blanket and a thermal probe. Eye ointment was applied to prevent dehydration. Before skin incision, mice were treated with buprenorphine (0.1 mg/kg i.p.) and lidocaine (0.4 mL/kg of a 1% solution, local application). After hair removal and disinfection with betadine and ethanol, the skin was opened, and the exposed cranial bone was cleaned and dried with cotton pads. Using a dental drill, the skull was thinned and a 5mm diameter bone was removed with the dura intact. For calcium imaging of cortical neurons, 200 nl of AAV1.syn.GCaMP6f.WPRE.SV40 (1×10^13^ vg/ml, University of Pennsylvania Vector Core) was injected at the following coordinates, V1:AP, -2.54 mm from bregma, L, +2.5; DV, -0.3 to - 0.1 mm and V2M: AP, -2.54 mm from bregma, L, +1.25; DV, -0.3 to - 0.1 mm from the skull using a Nanoject II^TM^ (Drummond Scientific) at a slow infusion rate (23 nL/s). For local deletion of CB1 receptors in layers 2/3 of V1 or V2M, CB1^floxed/floxed^ mice were injected with an AAV-CAG-Cre (0.5 μl) (Soria-Gomez et al., 2014) or a control virus AAV1.CAG.tdTomato.WPRE.SV40 (1.52×10^13^ GC/ml, University of Pennsylvania Vector Core). Animals were used for experiments five weeks after injection in order to have an optimal deletion of CB1, as previously described (Soria-Gomez et al., 2014;Soria-Gomez et al., 2015). After injection, the pipette was left in situ for an additional 5 min to prevent backflow. A circular cover glass (5 mm diameter) was placed over the exposed region and the glass edge was sealed to the skull with dental cement (Coffret SUPERBOND complete, Phymep). A circle stainless steel head post (Luigs & Neumann) was fixed to the mouse skull using dental cement.

### Habituation of the mice for awake imaging

Three weeks after the cranial surgery, mice were first habituated to the experimenter by handling. Mice were then accustomed to the imaging environment, where they freely moved on a rotating disk while being head-fixed. Habituation was performed for one week and mice did not receive any reward under any condition.

### *In vivo* two-photon calcium imaging

Imaging was performed using a two-photon microscope (Bruker Ultima) equipped with resonant galvo scan mirrors controlled by the Prairie software. Images were acquired using a water-immersion 20X objective (N20x-PFH, NA 1, Olympus). A 80-MHz Ti:Sapphire laser (Mai Tai DeepSee, Spectra-Physics) at 950nm was used for excitation of GCaMP6f and td-Tomato. Laser power did not exceed 30 mW under the objective. Fluorescence was detected by a GaAsP PMT (H7422PA-40 SEL, Hamamatsu). Time-series movies of neuronal populations expressing GCaMP6f were obtained at the frame rate of 30.257 Hz (294 x 294 μm, FOV; 0.574 μm/pixel). The duration of each focal plane movie was 330.5 seconds (10,000 frames). Animals were head-restrained and free to locomote on a disk treadmill (30 cm diameter) and kept under the two-photon microscope maximally for 70 minutes per day. The speed of each mouse on the rotating disk was continuously monitored using a rotary encoder (US Digital; H5-1024-IE-S; 1000 cpr) connected to a digitizer (Axon Digidata 1440, Molecular Devices) and analyzed with custom-written software in Python. Signals were recorded at 2KHz and downsampled to 30 Hz for analysis. For the calculation of speed, we multiplied the perimeter of the disk to the number of turns and then divided by the elapsed time.

To pharmacologically block CB1 we used the inverse agonist SR 141716A (Rimonabant). CB1-tdTomato mice were injected i.p. with Rimonabant (5 mg/kg, TOCRIS) or vehicle (1.25% (vol/vol) DMSO, 1.25% (vol/ vol) Tween80 in saline (Bellocchio et al., 2013) 30 minutes before the onset of the imaging session.

### Imaging data processing

Motion correction of the two-photon time-series was performed using a registration algorithm built into the Suite2p software (Stringer and Pachitariu, 2019). Segmentation into regions of interest (ROIs) was performed with Suite2p, confirmed by visual inspection and fluorescence traces extracted from the green channel for the different ROIs. Ca^2+^ traces were corrected for neuropil contamination. The neuropil mask resembled a band surrounding the ROI with its inner edge having a distance of 3 microns away from the edge of ROI and its outer edge having a distance of 30 microns from the edge of the ROI. The resulting neuropil trace, N, was subtracted from the calcium trace, F, using a correction factor a: Fc(t) = F(t) - a·N(t) where a was defined as 0.7 like previously described ((Stringer and Pachitariu, 2019). Changes in fluorescence were then quantified as ΔF/F_0_ where F_0_ was estimated as the average fluorescence value from a moving window of 60 seconds in order to remove slow time scale changes (Romano et al., 2017).

### Spike Deconvolution and spike tiling coefficient (STTC) calculation

STTC analysis was performed on deconvolved fluorescence traces as previously described (Cutts and Eglen, 2014). Briefly, a nonnegative spike deconvolution was applied to GCaMP6f fluorescence traces using the OASIS algorithm (implemented in Suite2P) with a fixed time scale of calcium indicator decay of 0.7s. Events corresponding to variations in fluorescence with an amplitude lower than 3 times the standard deviation (STDV) of the ΔF/F_0_ trace were excluded. STDV was calculated by fitting a Gaussian process to the negative ΔF/F_0_ fluctuations of each ROI (Romano et al., 2017). To quantify the correlation between event trains in pairs of ROIs (A and B), we look for events in A which fall within ±Dt of an event from B. This spike tiling coefficient reduces contribution of different firing rates in estimating correlations in neuronal spike times. The STTC value for sequences of calcium events in two ROIs, A and B, were calculated according to:

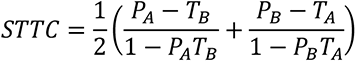

Where T_A_ is the proportion of total recording time which lies within ±Dt of any event in A. T_B_ was calculated in a similar manner. A Δt of 300 ms was used to take into account the slow rise time of GCaMP6f that limits precise temporal estimation of deconvolved events (Stringer and Pachitariu, 2019). P_A_ corresponds to the proportion of spikes from A which fall within Δt of any spike from B. P_B_ was calculated similarly. STTC values were calculated for all pairs of ROIs within the same individual field of view.

### Statistical Analysis

All statistical analysis were performed in Prism (GraphPad) or OriginPro (2016; OriginLab Co.). The normality of data was systematically tested with a D’Agostino and Pearson omnibus normality test. When normal, two datasets were compared using independent t-tests. When more than two data sets were compared, one-way ANOVAs and two-way ANOVAs were used. In situations where data was not normal or samples were small, we used non-parametric tests (Mann Whitney t-test and Kruskal-Wallis test followed by Dunn’s multiple comparison test for more than two groups, respectively) unless stated otherwise. For paired comparisons, we used paired *t-*test and Wilcoxon matched-pairs signed rank test for normally and non-normally distributed datasets, respectively. Differences were considered significant if p<0.05 (*p<0.05, **p<0.01, ***p<0.001) and means are always presented with the SEM.

## Acknowledgements

We thank Brahim Abbes for providing the Fiji plugin (Neurostack) for soma co-localization analysis and quantification. We would like to thank Ronan Chéreau and Michael Graupner for initial help in 2P- imaging implementation. We also thank members of the Bacci laboratory for helpful discussions. We also thank the ICM technical staff of the facilities PHENO-ICMICE, iGENSEQ and Histomics. All animal work was conducted at the PHENO-ICMice facility. We gratefully acknowledge Joanna Droesbeke for performing part of the transcardial perfusions. F.K. would like to thank the Fondation des Treilles for awarding her the 2020 Young Researchers Prize. This work was supported by “Investissements d’avenir” ANR-10-IAIHU-06; Agence Nationale de la Recherche (ANR-13-BSV4-0015-01; ANR-16-CE16- 0007-02; ANR-17-CE16-0026-01; ANR-18-CE16-0011-01; ANR-20-CE16-0011-01); Fondation Recherche Medicale (Equipes FRM – DEQ20150331684; Equipes FRM – EQU201903007860); DIM Region Ile de France, and a grant from the Institut du Cerveau et de la Moelle épinière (Paris) (A.B.); European Research Council (Starting Grant #678250) and the Brain & Behavior Research Foundation (NARSAD young investigator grant) (N.R.); the German Research Foundation through the Cluster of Excellence (EXC2067) Multiscale Imaging and the collaborative research center 889 (project B3) to OMS. The Core ICM facilities were supported by 2 “Investissements d’avenir” (ANR-10- IAIHU-06 and ANR-11-INBS-0011-NeurATRIS) and the “Fondation pour la Recherche Médicale”.

## Author Contributions

M.M., F.K., A.B. and J.L. designed the research; M.M., F.K., A.A., J.P., M.V., F.J.K., C.A., P.M. and J.L. collected data; M.M., F.K., A.A., J.P. and J.L., analyzed data; V.C., M.D.B.V.V., N.R. and A.B. provided analytical tools; G.M. and O.M.S. provided CB1 mouse lines and viral vectors; A.B. and J.L. co-supervised the project. M.M., F.K., A.B and J.L. wrote the manuscript. All authors edited the manuscript.

---

**Figure S1.**
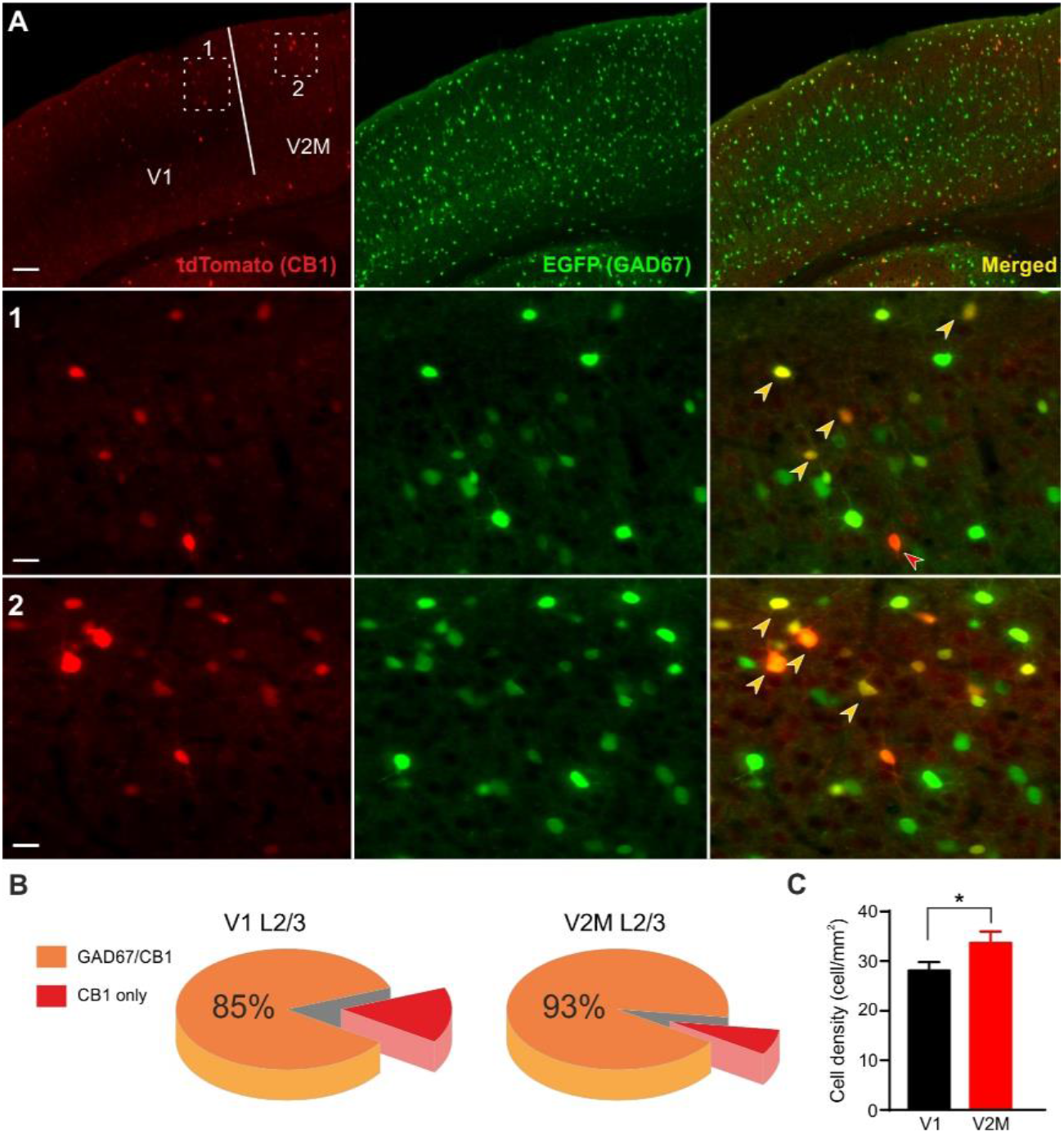
Density of CB1-expressing interneurons in V1 and V2M. **A:** Top row: Micrograph illustrating tdTomato (red) and EGFP (green) immunoreactivity in V1 and V2M in a CB1-tdTomato::GAD67EGFP mouse. This line was used to quantify CB1-expressing interneurons in L2/3 of the two visual areas. White dashed boxes are illustrated at increased resolution in the two rows below. Orange arrows illustrate CB1-expressing interneurons and red arrows illustrate CB1- expressing principal (pyramidal?) neurons. Scale bar: 100 (top row) and 25 µm (middle and bottom rows). **B:** Pie charts illustrating the fraction of CB1-expressing interneurons (co-localizing with EGFP) in the two visual areas. **C:** Cortical laminar density of CB1 INs in L2/3 of V1 and V2. Mean ± SEM, n=3 slices/animal, N = 5 animals for each condition. *: P < 0.05.s

**Figure S2.**
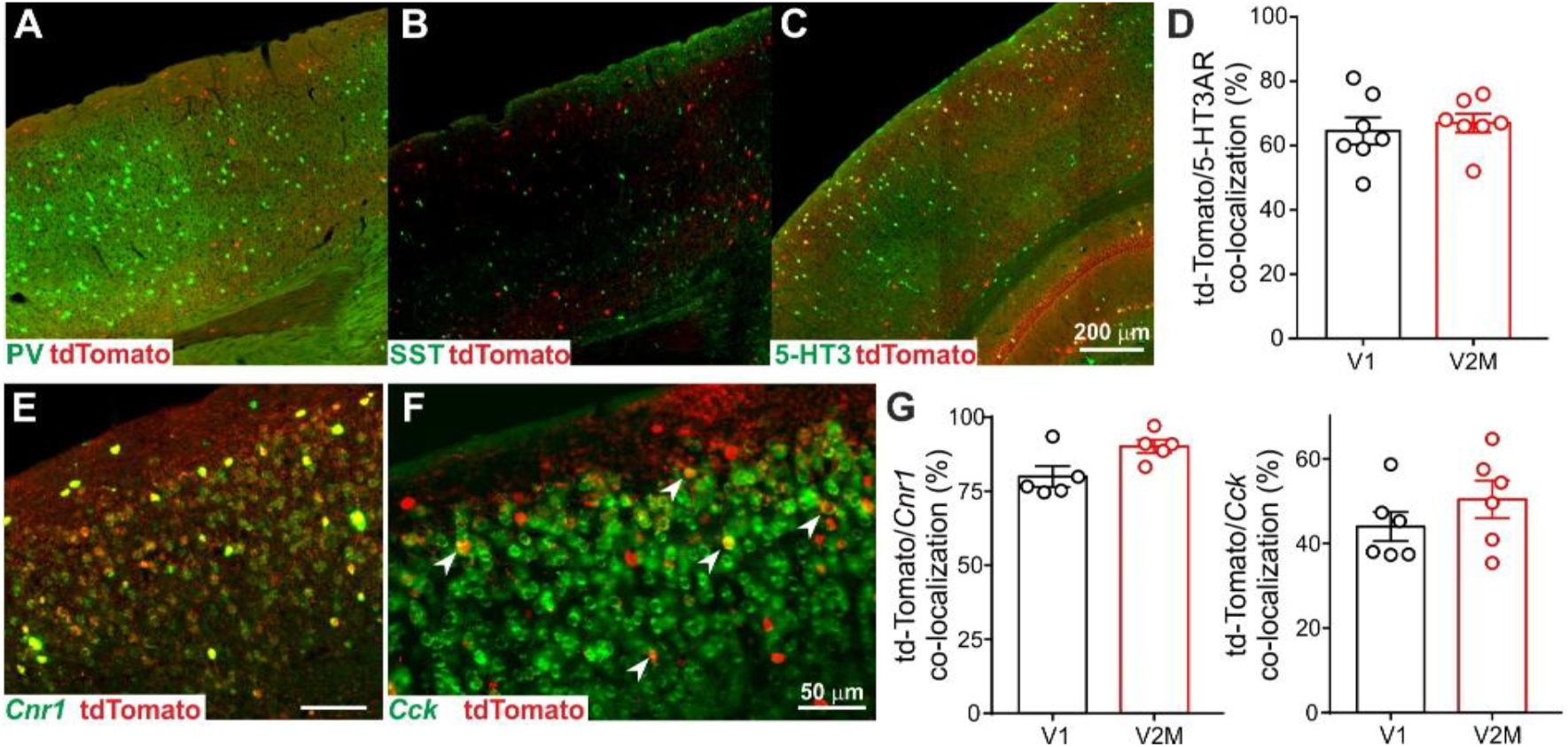
Lack of expression of PV and SST markers in CB1 BCs of the mouse visual cortex. **A:** Micrograph illustrating DsRed (red) and GFP (green) immunoreactivity against tdTomato and parvalbumin, respectively, in V1 and V2M. Note the lack of co-localization of parvalbumin marker with tdTomato CB1 BCs. **B:** Micrograph illustrating DsRed (red) and GFP (green) immunoreactivity against tdTomato and somatostatin, respectively, in V1 and V2M. Note the lack of co-localization of somatostatin marker with tdTomato CB1 BCs. **C:** Micrograph illustrating DsRed (red) and GFP (green) immunoreactivity in a 5-HT3-Cre::RCE mouse crossed with a CB1-tdTomato mouse. Therefore, GFP expression labels 5-HT3-expressing interneurons. Note extensive co-localization between the two markers. **D:** Summary plot of CB1/5-Ht3aR co-localization in V1 and V2M. **E:** Representative micrograph illustrating fluorescent *in situ* hybridization (FISH) against *Cnr1* (CB1) mRNA (green) and immunohistochemistry for DsRed in a CB1-tdTomato mouse to label td-Tomato-expressing neurons. **F:** Same as in E, but FISH was performed using anti-*Cck* probes. **G:** Summary graphs illustrating co- localization of *Cnr1* (CB1, left) and *Cck* (right) mRNA and td-Tomato. Please note the massive co- localization of CB1 mRNA with td-Tomato, validating the mouse line. Note also the extensive co-localization with CCK. For PV, n=3 slices/animal, N = 3 animals; for SST, = 4 slices/animals, N = 3 animals. For *Cnr1* n=4 slices/animal, N = 5 animals, for *Cck* n=6 slices/animal, N = 6 animals

**Figure S3.**
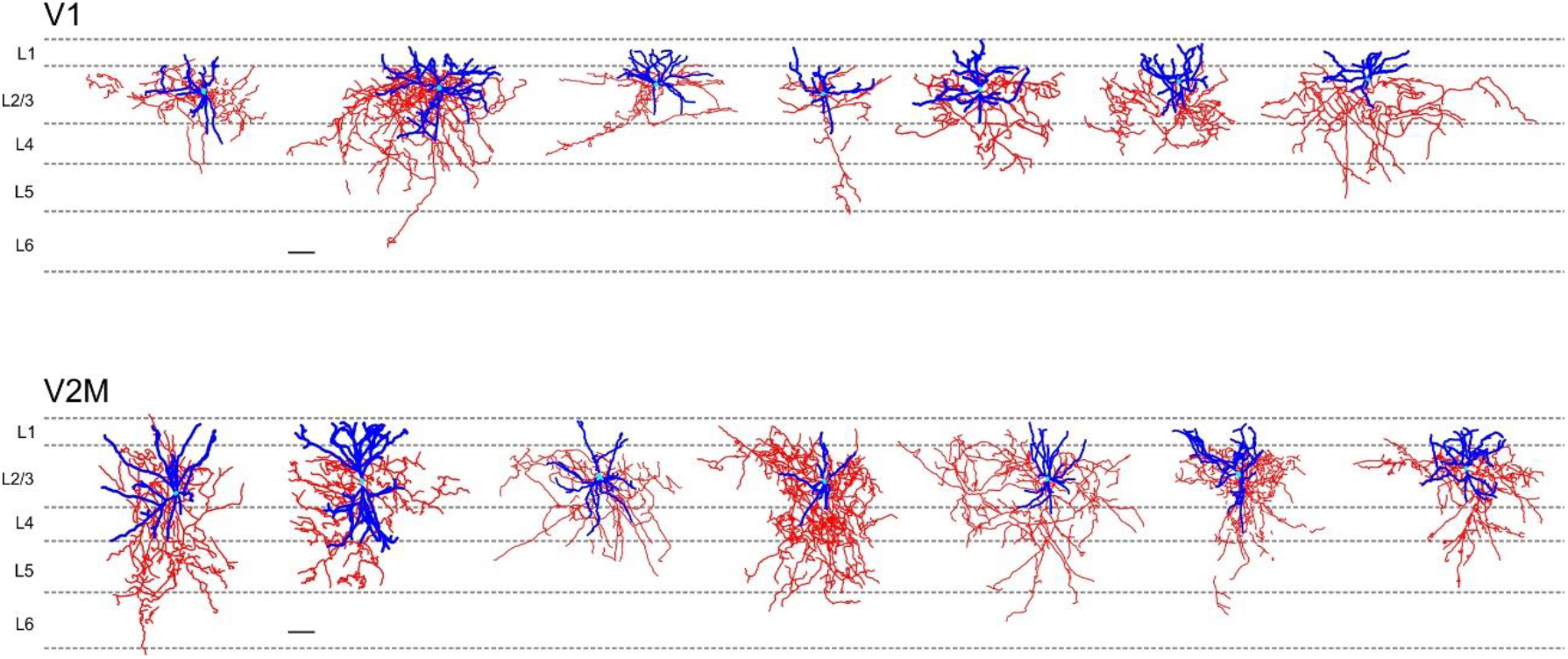
L4 of V2M is highly innervated by L2/3 CB1 BCs axons. **A:** Individual reconstructions of CB1 BCs in V1 filled with biocytin (5-8 mg/mL). Dendrites are in blue, axons in red, and soma in turquoise. **B:** Same as in A but for CB1 BCs in V2. Scale bars = 100 μm.

**Figure S4.**
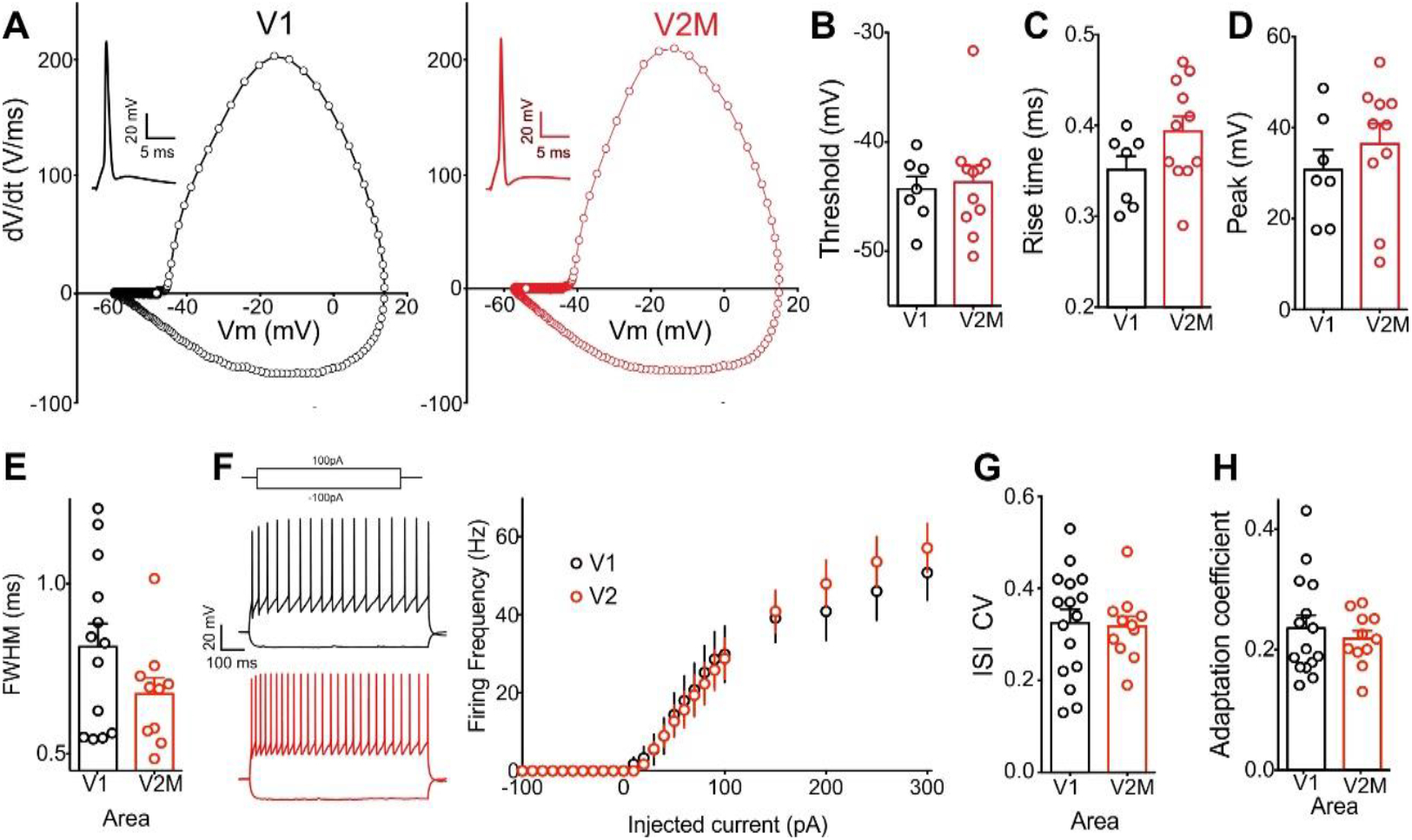
L2/3 CB1 INs exhibit similar active and passive single-cell properties in V1 and V2. **A:** Phase plane plots of the derivative of a single AP (insert) evoked by minimal current injection step (1.5 ms) in V1 (black) and V2 (red). **B-E**: Action potential parameters analyzed for characterization of the two CB1 BCs populations. **F:** Injection of negative and positive current in V1 (black trace) and V2 (red trace), with the resulting frequency/injected current curve (right). **G:** Coefficient of variation of the inter-spike interval of spikes elicited by depolarizing pulses. **H:** Adaptation coefficient corresponding to the ratio of the interval between the two first and two last spikes elicited by a depolarizing pulse.

**Figure S5.**
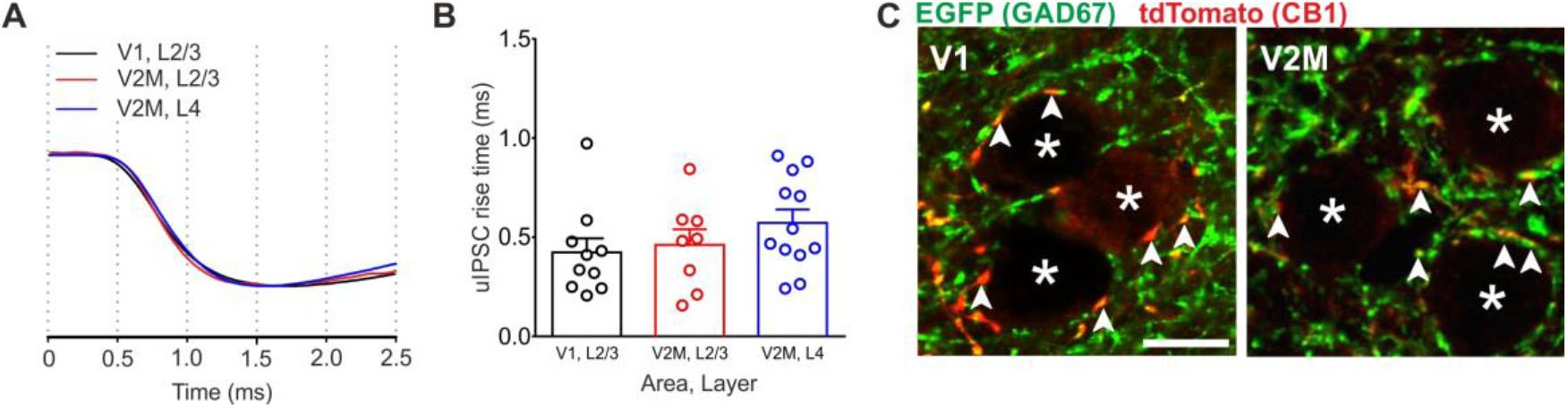
L2/3 CB1 INs from V1 and V2 exhibit perisomatic properties. **A:** Representative uIPSC traces illustrating fast rise time from L2/3 of V1 (black), L 2/3 of V2M (red) and L4 of V2M (blue). **B:** Population data the three synaptic connections displaying fast (<1ms) rise times. **C:** Micrograph illustrating DsRed (red) and GFP (green) immunoreactivity against tdTomato and GAD67, respectively, in V1 and V2M. Note the perisomatic pattern of innervation, typical of basket cells. Scale bars = 10 μm. Images were acquired with an inverted Confocal SP8 Leica DSL microscope.

**Figure S6.**
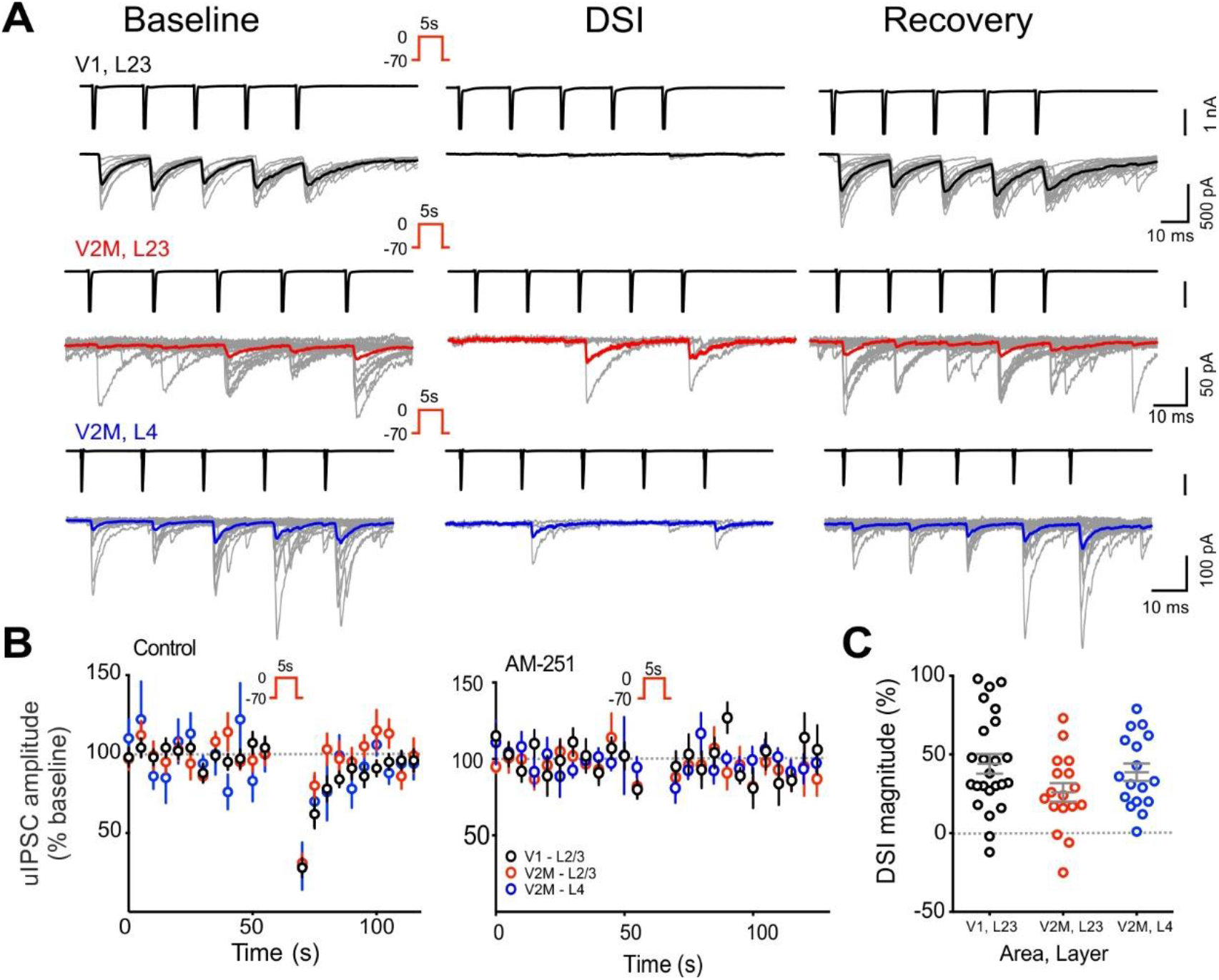
CB1-dependent DSI was robustly present in all three synapses. **A:** Representative traces in V1 L2/3 (top, black) V2 L2/3 (middle, red) and V2 L4 (bottom, blue) during the baseline (left), DSI (middle) and recovery from DSI (right). In all cases, presynaptic spikes above uIPSCs (black). Individual traces in grey and averages in the corresponding colour. Three sweeps were averaged for DSI and ten for baseline and recovery. Due to variability in uIPSC amplitude and high percentage of failures especially in V2M L4, the amplitude of postsynaptic responses to the presynaptic spike train was averaged in across the train (Vogel et al., 2016). B: Left panel, time course of average uIPSCs during baseline, DSI and recovery periods DSI in the three connections. The black dotted line represents the baseline value of 100 %. Right panel, time course of average uIPSCs during baseline, DSI and recovery periods DSI in presence of AM-251 for the three connections. **C:** Plot of individual average uIPSC amplitude after DSI. There is no differences in the maximum DSI values between connections in V1 L2/3, V2 L2/3 or V2 L4.

**Figure S7.**
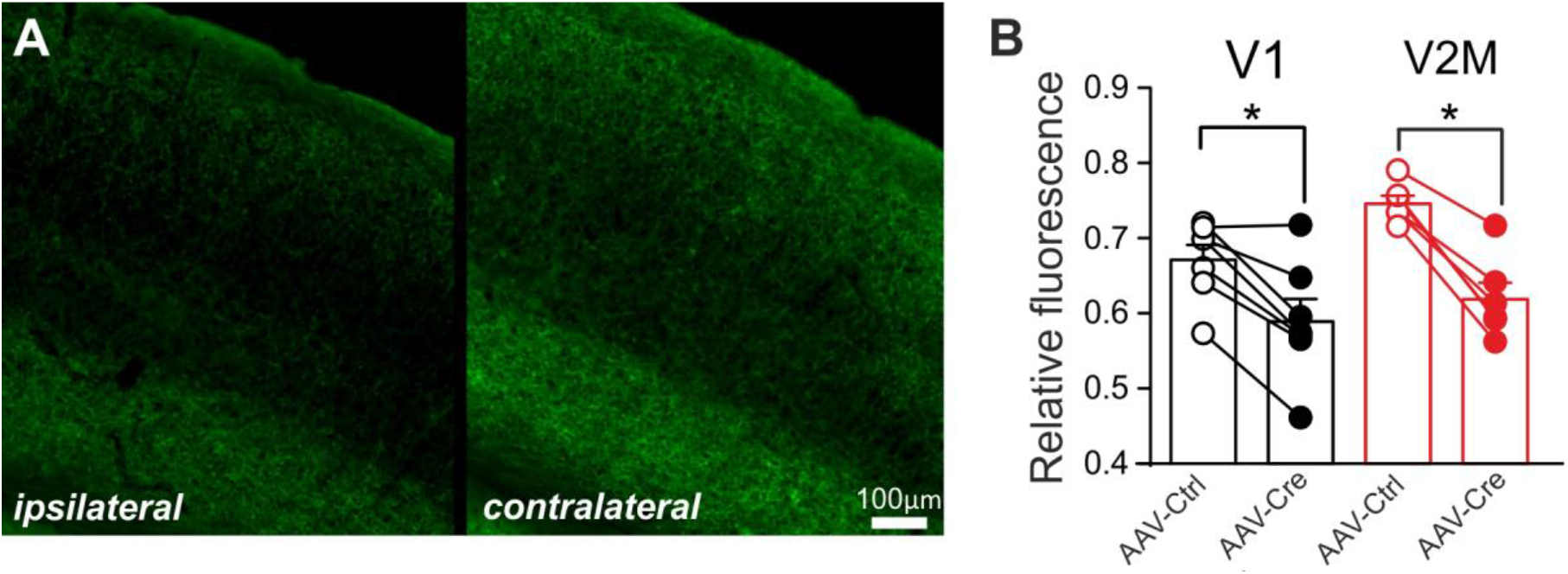
Quantification of CB1 expression after local AAV-Cre injection in CB1^floxed/floxed^ mice. **A:** Representative micrograph illustrating CB1 immunofluorescence in V1 of a CB1^floxed/floxed^ mouse injected with AAV-cre viral particles (ipsilateral, left) and its contralateral, non-injected site (right). Site of injection was identified by co-injection of AAV-tdTomato virus. CB1 pattern of expression was obtained using the “straight’ imageJ-option with a line width of 49 µm. **B:** Summary graph of CB1 immunofluorescence in AAV-control and AAV-Cre injected mice.

